# A chloride efflux transporter OsBIRG1 regulates grain size and salt tolerance in rice

**DOI:** 10.1101/2021.03.07.434240

**Authors:** Zhijie Ren, Fenglin Bai, Jingwen Xu, Li Wang, Xiaohan Wang, Qian Zhang, Changxin Feng, Qi Niu, Liying Zhang, Mengou Li, Jiali Song, Fang Bao, Liangyu Liu, Yikun He, Ligeng Ma, Jinlong Qiu, Wang Tian, Congcong Hou, Legong Li

## Abstract

Grain size is determined by the number of cells and cell size of the grain. Regulation of grain size is crucial for improving crop yield. However, the genes and underlying molecular mechanisms controlling grain size remain elusive. Here we report a member of Detoxification efflux carrier (DTX)/Multidrug and Toxic Compound Extrusion (MATE) family transporter, <u>BI</u>G <u>R</u>ICE <u>G</u>RAIN 1 (BIRG1), negatively regulates the grain size in rice. *BIRG1* is highly expressed in reproductive organs and roots. In *birg1* grain, the size of the outer parenchyma layer cells of spikelet hulls is noticeably larger but the cell number is not altered compared with that in the wild-type (WT) grain. When expressed in *Xenopus* oocytes, BIRG1 exhibits chloride efflux activity. In line with the role of BIRG1 in mediating chloride efflux, the *birg1* mutant shows reduced tolerance to salt stress under which the chloride level is toxic. Moreover, the *birg1* grains contain higher level of chloride compared to WT grains when grown under normal paddy field. The *birg1* roots accumulate more chloride than those of WT under saline condition. Collectively, our findings suggest that BIRG1 functions as a chloride efflux transporter regulating grain size and salt tolerance via controlling chloride homeostasis in rice.

## Introduction

Rice, as one of the most important crops in the world, provides the staple food for about half of the world’s population (Li *et al*., 2018). Thus, investigation of the genetic basis and molecular mechanism for rice grain yield regulation is of great significance. Grain size is an important yield trait, which is determined by grain width, length and thickness (Zuo and Li, 2014). In recent years, many quantitative trait loci (QTLs) and genes regulating grain size have been cloned and studied, providing insights into the molecular basis of the regulation of grain size (Cui et al., 2003; Huang et al., 2013; Ishimaru et al., 2013; Liu et al., 2015; Mao H, 2010; Shomura et al., 2008). However, additional genes that maintains ionic balance and controls this important trait remain to be identified.

Chloride (Cl^-^) is traditionally considered as a micronutrient (Broyer *et al*., 1954). It is involved in the stabilization of the photosystem II (PSII) and the regulation of enzyme activities such as the asparagine synthethase, amylases, and the tonoplast H^+^-ATPase (Metzler 1979; Rognes 1980; Churchill & Sze 1984; Kawakami *et al*., 2009). As a mobile anion in plant, Cl^-^ also plays main roles in the stabilization of the electric potential of cell membranes and the regulation of pH gradients (White & Broadley 2001; Hänsch & Mendel 2009). In addition, Cl^-^ serves as an osmotically active solute in certain tissues or single cells (e.g. guard cells) (Zonia *et al*., 2002; Hedrich 2012; Munemasa *et al*., 2015). On the other hand, Cl^-^ can be toxic to plants at high concentrations. The detrimental effects of high Cl^-^ concentration may be executed by interference with cell cycle regulation and inhibition of ribosomal enzymes that catalyze protein synthesis (Geilfus, 2018). Intriguingly, it has been reported that, during salt stress, the effects of Cl^-^ can be additive and/or synergistic to those of Na^+^. For instance, treatments with NaCl can affect the growth and physiology of rice more than treatments that contain only high concentrations of Cl^-^ or Na^+^ (Khare *et al*., 2015; Kumar & Khare 2015). Cl^-^ transport occurs primarily via the symplastic pathway (Brumós *et al*., 2010). Up to date, several plant transport proteins permeable to Cl^-^ have been identified and characterized, which includes AtCLCa (Chloride Channel) (de Angeli *et al*., 2006), AtCLCc (Jossier *et al*., 2010), AtCLCg (Nguyen *et al*., 2016), and AtALMT9 (ALuminium-Activated Malate Transporter) (Kovermann *et al*., 2007; de Angeli *et al*., 2013) in the shoot and AtNPF2.4 (Nitrate transporter /Peptide transporter Family) (Li *et al*., 2016), AtNPF2.5 (Li *et al*., 2017), AtSLAHs (SLow-type Anion Channel Associated/SLAC1 Homologues) (Cubero *et al*., 2016; Qiu *et al*., 2017), AtALMT9 (Baetz *et al*., 2016), AtCLCc (Jossier *et al*., 2010), CCCs (Cation Cl^−^ Cotransporter) (Zhang *et al*., 2010), AtNPF7.2, AtNPF7.3 (Li *et al*., 2017) and AtALMT12 (Meyer *et al*., 2010; Sasaki *et al*., 2010) in the root. Recently, it was reported that two tonoplast DTX/MATE proteins, DTX33 and DTX35, mediate chloride influx into the vacuole, which is essential for cell turgor regulation in *Arabidopsis* (Zhang *et al*., 2017).

DTX/ MATE transporters are conserved from bacteria to plants and animals (Brown et al., 1999; Hiroshi et al., 2006; Li et al., 2002). In *Arabidopsis*, several DTX/MATE members have been reported to function as transporters of organic acids and secondary metabolites (Li *et al*., 2002; Marinova *et al*., 2007; Yokosho *et al*., 2009; Zhang *et al*., 2014). For example, AtDTX1 mediates the efflux of plant-derived alkaloids, antibiotics, and other toxic compounds (Li *et al*., 2002). AtTT12 (TRANSPARENT TESTA 12) acts as a vacuolar flavonoid/H^+^-antiporter active in proanthocyanidin-accumulating cells of the seed coat (Marinova *et al*., 2007). The FRD3 (FERRIC REDUCTASE DEFECTIVE 3) mediates efflux of citrate into the root vasculature, which is necessary for efficient iron translocation (Green and L., 2004). The EDS5 (ENHANCED DISEASE SUSCEPTIBILITY 5) possibly functions as an SA (SALICYLIC ACID) transporter involved in the SA-dependent pathogen response pathway (Ding *et al*., 2014). Our previous work indicates that AtDTX50 may serve as an ABA (ABSCISIC ACID) efflux transporter (Zhang *et al*., 2014). RHC1 (RESISTANT TO HIGH CO_2_ 1) may act as a HCO_3_^-^ sensing component essential for CO_2_-induced stomatal closure (Tian *et al*., 2015).

The rice genome encodes about 45 members in the DTX/MATE family (Wang *et al*., 2016). Among them, three members (OsFRDL1 (FRD3-LIKE PROTEIN 1), OsFRDL2 and OsFRDL4) belong to the group of citrate transporters (Yokosho *et al*., 2009). OsFRDL1 is involved in the efficient translocation of Fe from the roots to the shoots (Yokosho *et al*., 2009). OsFRDL2 is involved in the Al-induced secretion of citrate (Kengo *et al*., 2016). OsFRDL4 is responsible for external detoxification of Al (Yokosho *et al*., 2016). The DTX/MATE family proteins transport a wide range of substrates and functions differently in various physiological processes. In this study, we report an uncharacterized member, <u>BI</u>G <u>R</u>ICE <u>G</u>RAIN 1 (BIRG1), functions as a chloride effluxer controlling chloride homeostasis, which negatively regulates grain size and positively contribute to salt tolerance in rice.

## Results

### Disruption of *BIRG1* increases grain size in rice

To explore the potential roles of DTX/MATE family genes in rice grain, we first analyzed the expression levels of the DTX/MATE family genes in spikes by searching the Rice Genome Annotation Project database (http://rice.plantbiology.msu.edu/). The results showed that 10 of the 46 *DTX/MATE* family genes are highly expressed in spikes (Table **S1**). Next, we individually mutated *Os06g29950*, *Os01g49120*, *Os02g57570*, *Os03g64150* and *Os08g43250* in the Nipponbare background using either TALENs (Transcription Activator-Like Effectors Nucleases) or CRISPR/Cas9 (Clustered Regularly Interspaced Short Palindromic Repeats-CRISPR-Associated protein 9) technologies (Fig. **S1a**). We found that a TALENs-mutagenized mutant harboring 7-base deletion at the ORF of *Os03g64150*, showed increased grain size (**Fig. 1a, b**). We named it as *<u>B</u>IG <u>RI</u>CE <u>G</u>RAIN 1* (*BIRG1*). To further test if the increased grain size is caused by the loss-of-function mutation of the *BIRG1*, we generated an independent *BIRG1* knockout line, *birg1-2* (harboring 1-base deletion at the target site that causes a frame shift), using CRISPR/Cas9 technology (Fig. **1a**). Indeed, compared with the WT, *birg1-1* and *birg1-2* both had significantly increased grain length, width and thickness (Fig. **1e, f**; Fig. **S1f, g, h, i**). As a result, the 1,000-grain weight was increased by 13.03% and 13.44% in *birg1-1* and *birg1-2*, respectively (Fig. **1d**; Fig. **S1c, d, e**). When plants were grown naturally in three different paddy fields where the Cl^-^ level varied but is not detrimental, the increased grain size phenotype of *birg1* mutant was observed consistently (Fig. **S2**). This result indicates that BIRG1 protein regulates rice grain size under a wide range of external Cl^-^ level. Moreover, we generated *BIRG1* knockdown mutants using RNA interference (RNAi) technology. Similar to *birg1-1* and *birg1-2*, the *BIRG1* RNAi lines, *Ri-1* and *Ri-2* also showed increased grain length and width (Fig. **S3**). Taken together, these data indicate that *BIRG1* is a negative regulator of the grain size in rice.

**Fig. 1.**
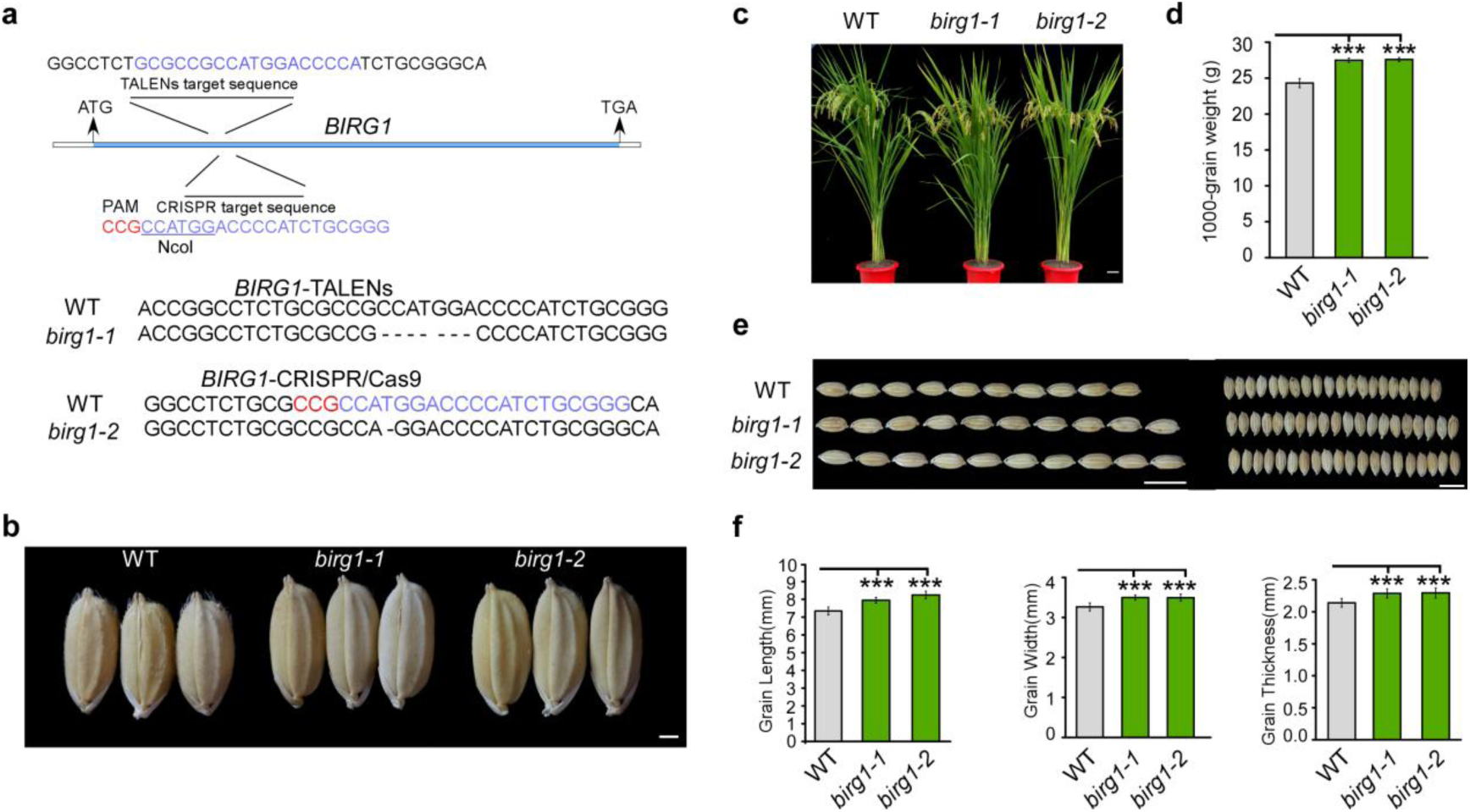
Knockout of *BIRG1* increases grain size in rice. (a) Characterization of *birg1-1* and *birg1-2* generated by TALENs and CRISPR/Cas9 techniques, respectively. (b) Grain morphology of the WT, *birg1-1* and *birg1-2*. (Scale bar=1 mm). (c) Gross morphology of 120 days-old plants of WT, *birg1-1* and *birg1-2* grown in a paddy field. (Scale bar=4 cm). (d) Statistical data of 1,000-grain weight in the WT, *birg1-1* and *birg1-2*. (n=3). (e) Comparative observation on grain arrangement of the WT, *birg1-1* and *birg1-2* (Scale bar=1 cm). (f) Statistical data of the grain length, grain width, and grain thickness in the WT, *birg1-1* and *birg1-2*. Rice plants were grown in a paddy field located in Lang Fang, He Bei, China, where the Cl- concentration is 0.11±0.06 g kg^-1^, under natural conditions. Means ± SD are given in F (n =10), *p* value (Student’s *t*-test), * *p*<0.05, ** *p* <0.01 and *** *p* <0.001.

Compared with the WT, the *birg1-1* and *birg1-2* mutants showed negligible difference in the gross morphology from the seedling stage to vegetative developmental stage (Fig. **S1j, k**). However, at mature stage, plant height of *birg1* mutants was slightly decreased which attributed to reduced length of the uppermost internode (Fig. **1c**; Fig. **S1l**; Fig. **S4a, b**). The architecture of the *birg1* mutants was more upright, short and compact (Fig. **1c**), which endowed the *birg1* mutants an increased lodging resistance (Fig. **S5**). Moreover, the *birg1* mutants displayed noticeable chlorosis at the leaf tips and edges (Fig. **1c**). Compared with the WT, there was no difference at flag leaf length and width, tiller number and grain density of per panicle in the *birg1* (Fig. **S4c, d, e, h**). However, the panicle length of the *birg1* was shorter than that of the WT (Fig. **S4f**). Grain number per panicle and seed setting rate of *birg1* were reduced compared to those of the WT (Fig. **S4g, i**). These results suggest that *BIRG1* may influence the balance between grain number and grain size. The *birg1* mutant displayed pleotropic phenotypes as described. We postulated that some of the phenotypes, such as decreased plant height, reduced grain number and lower seed setting rate, may attribute to compromised photosynthetic capacity of the *birg1* mutant. To test this possibility, we measured the photosynthetic rate of *birg1* and WT. Our results showed that, indeed, the photosynthetic rate of *birg1* is significantly lower than that of the WT (Fig.**S6**).

### Expression level of *BIRG1* is induced in reproductive organs and its protein localizes at the plasma membrane

We next performed reverse transcription quantitative PCR (RT-qPCR) to investigate the spatial expression profile of *BIRG1*. The RT-qPCR results revealed that *BIRG1* was expressed in the root, culm, leaf, sheath, anther, and panicle, with the relatively higher expression levels in the root and anther (**Fig. S7a**). Importantly, during the earlier stage of panicle development, the expression level of *BIRG1* increased significantly with panicle elongation (**Fig. 2a**). In addition, we cloned a 3.2-kb promoter region of *BIRG1* and generated the promoter-*GUS* transgenic rice plants. GUS staining showed that *BIRG1* was expressed abundantly in the root, culm node, grain, leaf and guard cells (**Fig. 2b**), which is consistent with the results from RT-qPCR analysis. In the panicles, strong GUS staining was also observed in spikelet hulls, pistil, anther, pollens and seeds (**Fig. 2c**). The expression level is induced in reproductive organs, not in other tissues, indicating *BIRG1* might be involved in the grain filling.

**Fig. 2.**
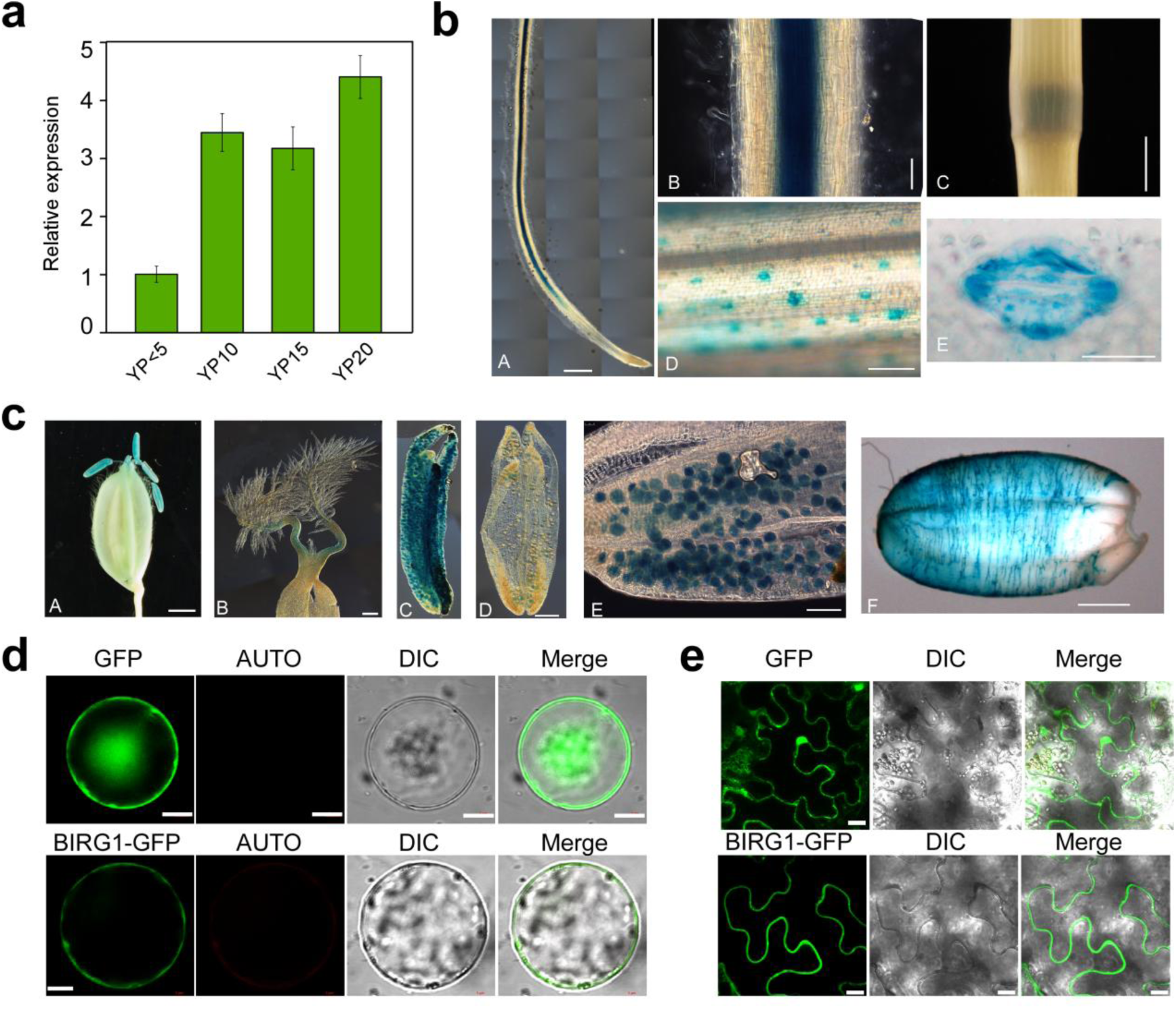
Tissue-specific expression and subcellular localization of *BIRG1*. (a) Expression of *BIRG1* in different lengths of developing young panicles (indicated as numbers, cm). (b) Expression of *BIRG1*::GUS in young roots (A and B) (Scale bar, a=0.1 cm, b=0.05 cm), culm node (C) (Scale bar, 1 cm), leaf (D) (Scale bar, 1 cm) and guard cells (E) (Scale bar, 8 μm). (c) Expression of BIRG1::GUS in spikelet hulls (A) (Scale bar, 0.2 cm), pistil (B) (Scale bar, 0.5 mm), anther (C and D) (Scale bar, 1 mm), pollens (E) (Scale bar, 1 mm) and seed (F) (Scale bar, 1 mm). (d) Rice green tissue protoplasts transformed with 35S:: *BIRG1*-*GFP* and 35S::*GFP*. Scale bar =5 μM. (e) Tobacco epidermal leaf cells transiently transformed with 35S:: *BIRG1*-*GFP* and 35S::*GFP*. Scale bar =50 μm.

Previous study has shown that several members of DTX/MATE family protein, for examples DTX35 and DTX33, are localized to tonoplast (Zhang *et al*., 2017). As the subcellular localization of a transporter is critical for its function, we first examined the subcellular localization of BIRG1 by the GFP fusion approach. We made *BIRG1::GFP* fusion constructs under the control of *35S* promoter and transiently expressed them in rice protoplasts and tobacco (*Nicotiana benthamiana*) epidermal cells, respectively. In rice protoplast, the green fluorescence of BIRG1-GFP was observed at the protoplast plasma membrane compared with the diffuse cytoplasmic localization of the eGFP control (**Fig. 2d**). Consistently, in tobacco epidermal cells, the BIRG1-GFP was targeted at the plasma membrane, whereas the eGFP control was distributed in the nucleus and plasma membrane (**Fig. 2e**). These results indicated that BIRG1 is localized to the plasma membrane.

### The *birg1* mutant shows accelerated grain filling rate

Given the importance of grain filling in contributing to the seed size, we investigated the grain filling rate by measuring fresh and dry weight of the grains and brown rice during grain filling (Fig. **3a**). All fresh and dry weight of grains and brown rice of *birg1-1* were higher than those of the WT starting from ∼12 days after fertilization (DAF). The differences became more significant along with the grain development. The maximum grain filling rate was achieved at ∼21 DAFs. At this point, fresh and dry weight of the grains of *birg1-1* were 18.1% and 21.1% higher than the WT; Fresh and dry weight of the brown rice of *birg1-1* were 16.0% and 18.2% higher than the WT, respectively (Fig. **3b, c, d, e**). These data indicates that the grain filling rate of *birg1* is accelerated compared to the WT. As the grain filling is actually a process of starch accumulation, we examined the expression levels of five genes related to starch synthesis *RSR1/AP2/EREBP* (*RICE STARCH REGULATOR 1/AP2/EREBP FAMILY TRANSCRIPTION FACTOR*), *AGPL2* (*ADP-GLUCOSE PYROPHOSPHORYLASE LARGE SUBUNIT 2*), *GBSSI* (*GRANULE-BOUND STARCH SYNTHASE 1*), *GLUCAN* (*GLUCAN SYNTHASE*), *BEI* (*BRANCHING ENZYME I*) (Wang *et al*., 2015). Our result showed that all these five genes were up-regulated to different extent in the *birg1* mutants, which is correlated with the faster grain filling rate of *birg1* (Fig. **S7b**).

**Fig. 3.**
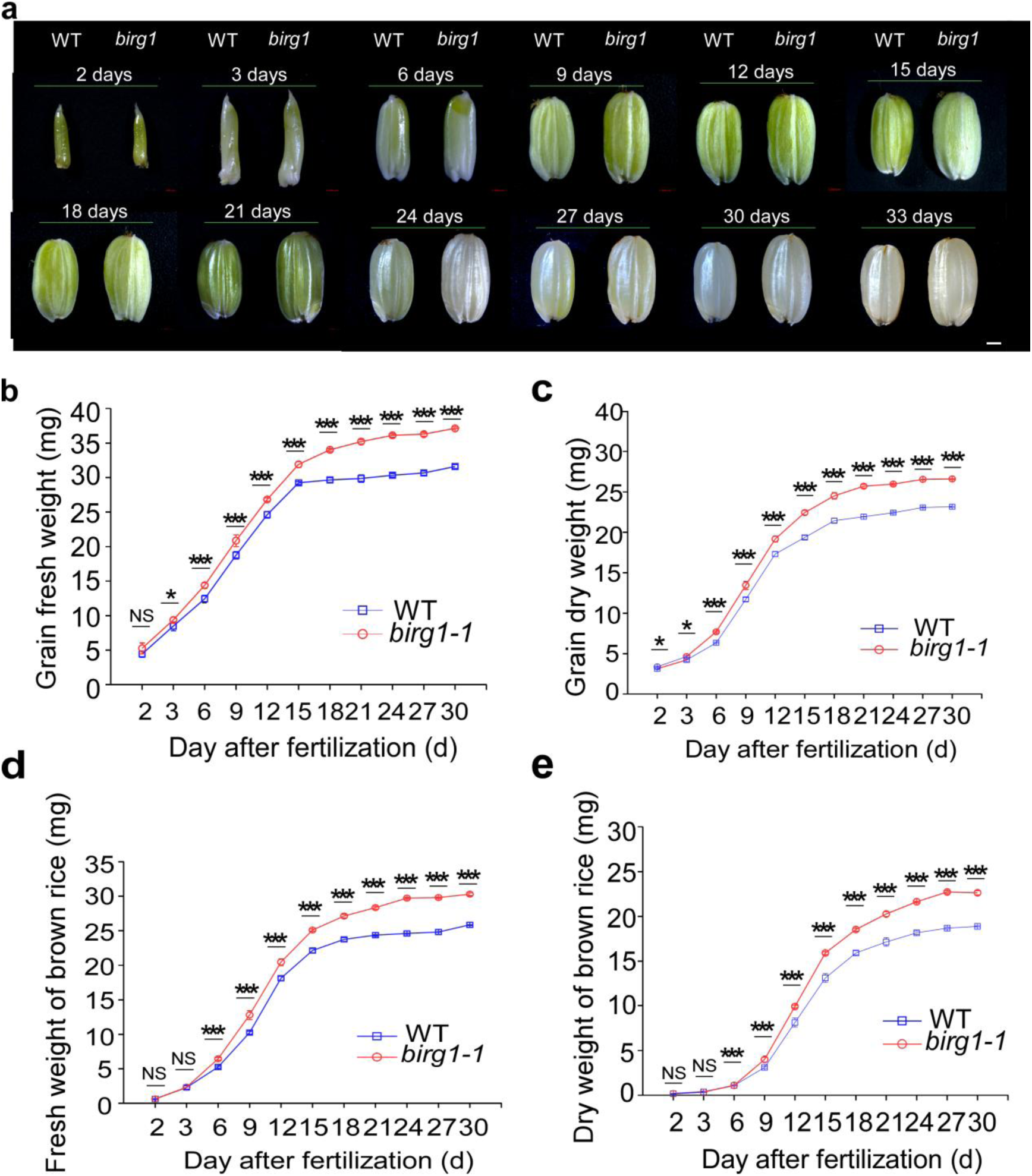
The *birg1* mutant shows accelerated grain filling rate. (a) Developing WT and *birg1-1* seeds in 2, 3, 6, 9, 12, 15, 18, 21, 24, 27, 30 and 33 DAP were photographed. Bars =1 mm. (b) Time-course of the WT and *birg1-1* grain fresh weight (n =100 grains for each point). (c) Time-course of the WT and *birg1-1* grain dry weight (n =100 grains for each point). (d) Time-course of the WT and *birg1-1* brown rice fresh weight (n =100 grains for each point). (e) Time-course of the WT and *birg1-1* brown rice dry weight (n =100 grains for each point). WT, Line with blue diamonds; *birg1-1*, Line with red circle. Means ± SD are given in B-E (n =5*20), *P* Value based on Student’s *t*-test. NS means No Significance, * *p*<0.05, ** *p* <0.01 and *** *p* <0.001.

On the basis of the observations that the filling rate of the *birg1* mutant is accelerated and the *birg1* matures earlier than the WT (Fig. **3a**), we asked whether the starch accumulation in the endosperm of the *birg1* mutant is affected. We found that the mutant had a loose endosperm texture and loose, small starch granules, whereas the WT starch accumulated relatively densely (Fig. **S8a**), indicating that the grouting of the mutant stopped before it was fully completed. In addition, there was no significant difference in crude protein content and moisture content between the grains of WT and *birg1* (Fig. **S8b, c**). The amylose content in the *birg1* grain was significantly higher than that in the WT (Fig. **S8d**). At the same time, the viscosity of *birg1* pasting starch is higher than that of the WT (Fig. **S8e**), which may make the *birg1* rice taste better.

### Cytological alterations in *birg1* mutant grain

The *birg1* mutants showed increased grain size. It has been proposed that grain size is determined by the size of spikelet hull, which consists of a lemma and a palea (Na and Li, 2015). Changes in both cell size and cell number affect the final size of a spikelet hull (Na and Li, 2015). To uncover the cellular mechanism for BIRG1 in grain size, we investigated cells in WT and *birg1* spikelet hulls using a scanning electron microscope. We found that, in *birg1-1* and *birg1-2*, the lemma cell length and width were greater than those in the WT (Fig. **4a, b**). The palea cell length in *birg1-1* and *birg1-2* mutants were greater than that in the WT (Fig. **4a, c**), but there was no significant difference in cell width of the palea between the mutants and the WT (Fig. **4c**).

**Fig. 4.**
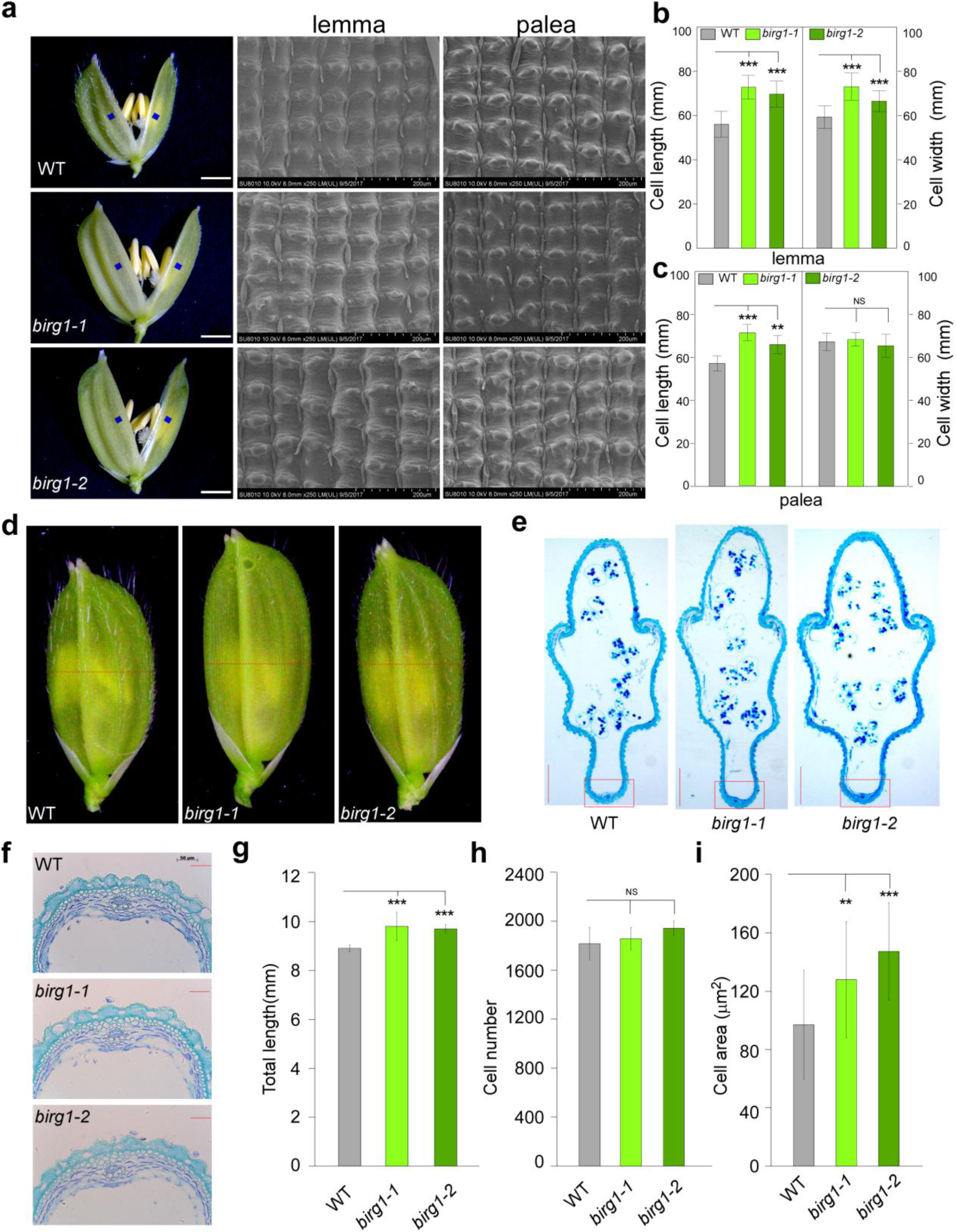
Increased grain size in *birg1* mutants is caused by increased cell expansion in the spikelet hulls. (a) Scanning electron and light microscope photographs of the outer surfaces of young spikelet hulls of WT, *birg1-1* and *birg1-2*. Scale bar, 1 mm for the whole spikelets and 200 µm for lemma and palea. (b) Comparison analysis of cell length and width in outer surfaces of lemma (n=20). Scale bar=500 μm. (c) Comparison analysis of cell length and width in outer surfaces of palea (n=20). Scale bar=100 μm. (d) Young spikelet hulls of WT, *birg1-1* and *birg1-2*. The red line indicates the position of the cross-section. (e) Cross-sections of spikelet hulls. Scale bar=0.5 mm. (f) Magnified view of the cross-section area boxed in (e). Scale bar=50 μm. (g) Comparison analysis of the total cell length in the outer parenchyma layer (n = 5). (h) Comparison analysis of the total cell number in the outer parenchyma layer (n = 5). (i) Comparison analysis of the cell area in the outer parenchyma layer (n = 5). All data are means ± SD. Student’s *t*-test was used to generate the *P* values; * *p*<0.05, ** *p* <0.01 and *** *p* <0.001.

Next, histological analysis was performed on the spike hulls two days before pollination (Fig. **4d, e**). The results showed that the outer parenchymal cells of the *birg1-1* and *birg1-2* mutants were larger than those of the WT (Fig. **4f, i**), thereby increasing the total length of the outer parenchymal cells (Fig. **4f, 4g**). Further statistical analysis on the number of outer parenchyma cells found no significant difference between WT and mutants (Fig. **4h**). In support of these results, expression levels of five cell cycle-related genes *CDKA1* (*CYCLIN-DEPENDENT KINASE A-1*), *CYCD1;1* (*CYCLIN -D1-1*), *CYCT1* (*CYCLIN -T1*), *H1* (*CYCLIN -H1*) and *E2F2* (*E2F TRANSCRIPTION FACTOR*) in the WT and *birg1* were comparable, whereas, among the five cell expansion-related genes tested (*EXA5*, *EXA10*, *EXB3*, *EXPB4* and *EXB7*), *EXB3* and *EXB7* were significantly elevated in young panicles of the *birg1* mutant (Fig. **S4c, d**) (Li *et al*., 2011; Liu *et al*., 2015). Together, these results demonstrated that the larger spikelet in *birg1* was caused by the enlargement of the single cell but not the change in cell number, suggesting that *BIRG1* regulates spikelet size by affecting cell expansion.

### BIRG1 can mediate chloride efflux

The genetic evidences above demonstrated that loss-of-function of *BRIG1* resulted in expanded cell size and increased grain size. As we know, BIRG1 is a member of the MATE family proteins that are found in bacteria, fungi, plants, and animals (Brown *et al*., 1999; Hiroshi *et al*., 2006; Li *et al*., 2002). Studies have shown that MATE proteins transport various substrates. Among the plant MATE-type transporters, some PM-focused members select citric acids as substrates and some tonoplast localized members function as anion channels that mediate transport of halogen ion (Magalhaes *et al*., 2007; Maron *et al*., 2009; Zhang *et al*., 2017).

To investigate the electrophysiological properties of BIRG1, we expressed the BIRG1 protein in the *Xenopus laevis* oocytes and examined its activity using various ionic buffers. Two-electrode voltage clamp (TEVC) recording showed that the oocytes injected with *BIRG1* cRNA produced large inward currents of ∼2200 nA, at -140 mV in bath solution containing 48 mM NaCl, which were not observed in the control cells (water injected) (Fig. **5a**). By convention, inward currents reflect positive charge influx or negative charge efflux. When we replaced Cl^-^ in the bath solution with gluconate (an impermeable anion), the large inward currents disappeared in BIRG1-expressing cells. These results indicate that the inward currents may be produced by chloride efflux (Fig. **5a, b**). Upon changing the bath Cl^-^ concentration, the magnitude of the inward current changed proportionally (Fig. **5c, d**), indicating that the BIRG1-mediated inward currents were dependent on external Cl^-^. In addition, we applied a typical anion channel blocker 4,4’-diisothiocyanostilbene-2,2’-disulfonic acid (DIDS) to the bath solution and observed inhibition of the BIRG1 currents (Fig. **5e, f**). We also examined the anion selectivity of BIRG1 by perfusing the bath solution with various anions. The results showed a selectivity sequence of Br^-^ > Cl^-^ > NO_3_^-^ ≈ SO_4_^2-^ > malate^2-^ > gluconate^-^ (Fig. **5a, b**). Next, to further clarify the rectification property of BIRG1, we removed extracellular Cl^-^ by using a gluconate-based bath medium and utilized the Cl^-^ releasing from the electrodes as the charge carrier. The method was used successfully for recording the channel activity of SKOR, an outward potassium channel from *Arabidopsis* (Gaymard et al., 1998; Li et al., 2008a). Intriguingly, we found that, similar to what observed in SKOR-expressing oocytes, the BIRG1-expressing oocytes only produced small inward currents at the time point of 1 minute (Fig. **5g, h, i, j**). However, as time goes by, the inward currents became remarkably larger in the same oocyte, suggesting increased Cl^-^ efflux across the plasma membrane (Fig. **5g, h, i, j**). No inward current was observed in water-injected control oocytes (Fig. **5g, h, i, j**). Taken together, these data confirmed that BIRG1 can mediate anion efflux, especially inorganic anions, such as Cl^-^.

**Fig. 5.**
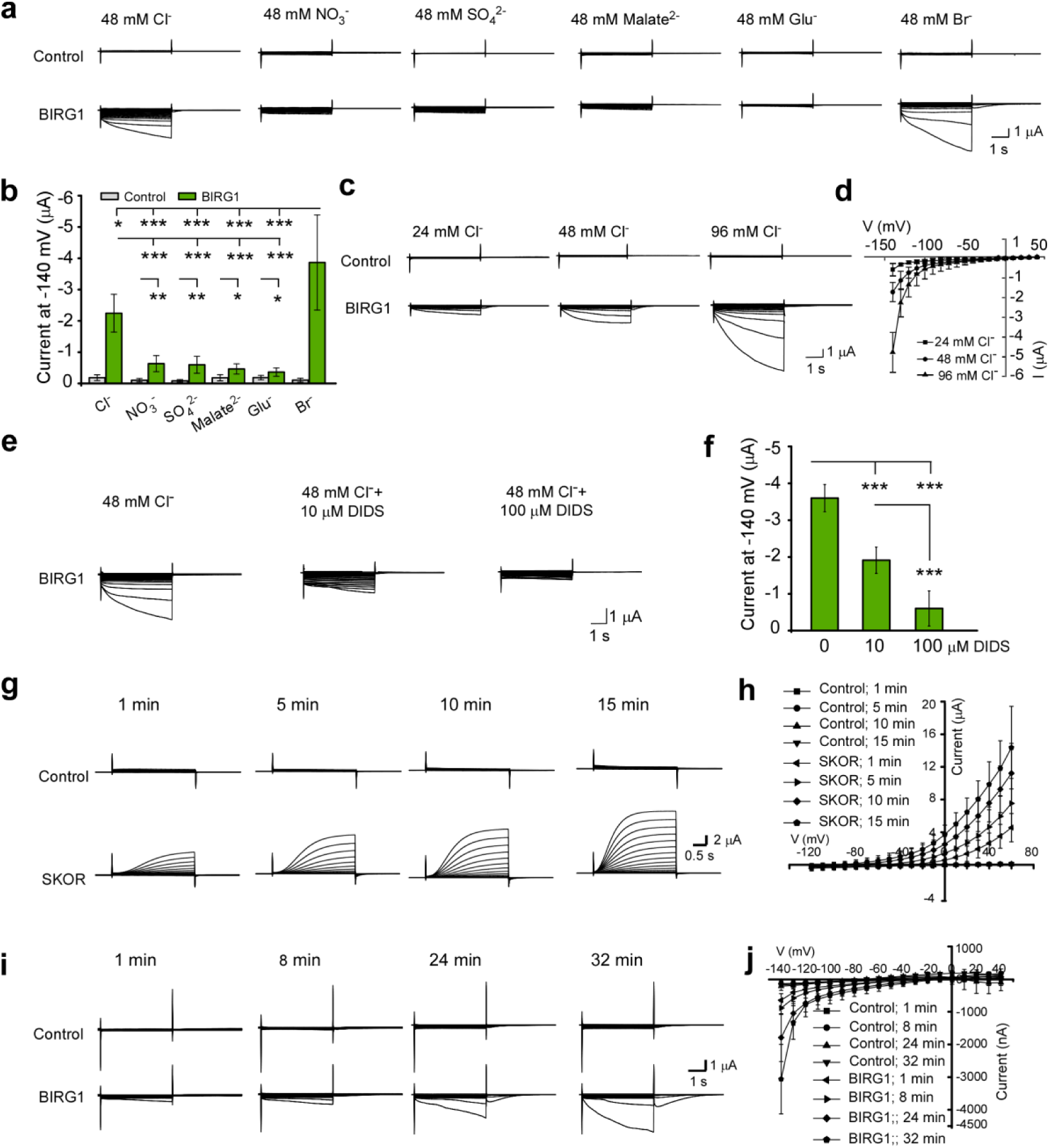
BIRG1 mediates Cl^-^ efflux in *Xenopus* ooctyes. (a) Typical whole-cell currents recorded by hyperpolarized pulses from oocytes injected with water (“Control”) or oocytes expressing BIRG1 perfused with a bath solution containing 48 mM Cl^-^, NO_3_^-^, SO_4_^2-^, Malate^2-^, Br^-^ or Glu^-^. Voltage ramps of 4 s duration were applied to the membrane with a voltage ranging from -140 to +40 mV. (b) Current amplitudes at −140 mV from multiple recordings as in (a). (c) The typical current traces generated by control oocytes injected with water or oocytes expressing BIRG1 bathed in a solution containing 24, 48 or 96 mM chloride with a voltage ranging from -140 to +40 mV. (d) The current–voltage relationship was deduced as in (c). The data are mean ± SD (n = 5). (e) Oocytes expressing BIRG1 were recorded in a solution containing 48 mM Cl^-^ plus 0, 10 or 100 μM DIDS. Voltage ranging from -140 to +40 mV. (f) Current amplitudes at -140 mV in the presence of indicated concentrations of DIDS as in (e). (g) Typical whole-cell currents recorded from control oocytes or oocytes expressing SKOR perfused with standard ND96 solution. The currents were recorded at the time point indicated. Voltage ranging from -120 to +60 mV in 10 mV increments. The holding potential was -100 mV. (h) The current–voltage relationship was deduced as in (g). (i) Typical whole-cell currents recorded from control oocytes or oocytes expressing BIRG1 perfused with a modified Cl^-^-free ND96 (chloride was replaced with glutamate). The currents were recorded at the time point indicated. Voltage ranging from -140 to +40 mV in 10 mV increments. (j) The current–voltage relationship was deduced as in (i). All data are mean ± SD (n≥5), *p* Value based on Student’s *t*-test, * *p*<0.05, ** *p* <0.01 and *** *p* <0.001.

### BIRG1 contributes to salt tolerance in rice

Under high salinity, Cl^-^ usually accumulates to toxic level in the cytoplasm. Our results demonstrated that BIRG1 is expressed highly in the roots and it exhibits chloride efflux activity in oocytes. We speculated that BIRG1 may expel chloride from the cytosol to the apoplast to reduce the overall chloride content in the plant thus playing a role in salt tolerance. To test this possibility, the WT and *birg1* seeds were geminated and grown in half strength MS medium containing 100 mM or 150 mM NaCl for 14 days. Clearly, the growth of WT plants was severely inhibited by the presence of 100 mM or 150 mM NaCl (Fig. **6a**). Intriguingly, the inhibition effect of NaCl on *birg1* mutants was more pronounce, as indicated by the observation that the roots and shoots of *birg1* mutants are shorter than those of WT under the salt stress conditions (Fig. **6b, c, d, e, f, g**). It is well known that high salinity imposes both ionic and osmotic stresses to plants. To distinguish if the hypersensitivity of *birg1* to NaCl is caused by its osmotic effect, the WT and *birg1* seeds were also subjected to 300 mM mannitol for 14 days. Our results showed that the growth of WT and *birg1* mutants were greatly suppressed to similar degree (Fig. **6h, i**), indicating that *BIRG1* is not involved in hyper-osmolality stress. Moreover, we found that when grown in the medium containing 150 mM NaNO_3_, the WT and *birg1* showed no morphological difference (Fig. **6h, i**), suggesting that BIRG1 does not respond to NO_3_^-^, which is in line with the result from TEVC experiment that BIRG1 exhibits very low permeability to NO_3_^-^ (Fig. **5a, b**). Together, these results demonstrated that BIRG1-mediated chloride efflux contributes to salt tolerance in rice.

**Fig. 6.**
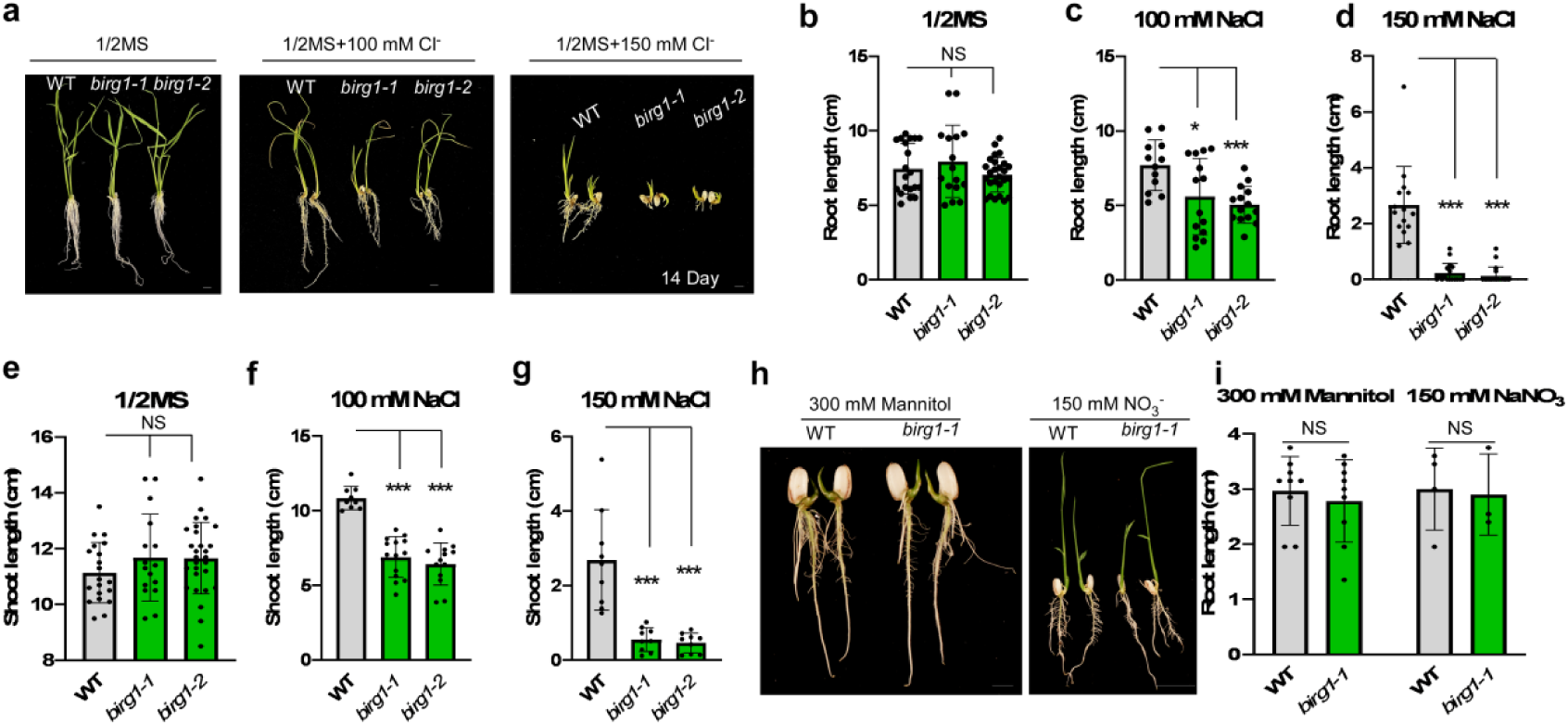
NaCl stress assays for seedlings in WT and *birg1*. (a) Seedling morphology of the WT, *birg1-1*, and *birg1-2* grown on 1/2MS, 1/2MS+100 mM NaCl and 1/2MS+150 mM NaCl for 14 days. (Scale bar=1 cm). (b) The root length of WT, *birg1-1*, and *birg1-2* grown on 1/2MS, n≥15. (c) The root length of WT, *birg1-1*, and *birg1-2* grown on 1/2MS+100 mM NaCl, n≥15. (d) The root length of WT, *birg1-1*, and *birg1-2* grown on 1/2MS+150 mM NaCl, n≥15. (e) The shoot length of WT, *birg1-1*, and *birg1-2* grown on 1/2MS, n≥15. (f) The shoot length of WT, *birg1-1*, and *birg1-2* grown on 1/2MS+100 mM NaCl, n≥15. (g) The shoot length of WT, *birg1-1*, and *birg1-2* grown on 1/2MS+150 mM NaCl, n>9. (h) Seedling morphology of the WT, *birg1*grown on 1/2MS+300 mM Mannitol, and 1/2MS+150 mM NaNO_3_. (Left Scale bar=1 cm and right Scale bar=2 cm). (i) The root length of WT, *birg1* grown on 1/2MS+300 mM Mannitol, and 1/2MS+150 mM NaNO_3_. n>3. The data are mean ± SD, Student’s *t*-test was used to generate the *P* values, * *p*<0.05, ** *p* <0.01 and *** *p* <0.001.

### BIRG1 regulates chloride homeostasis in rice

Cl^-^ is one of the essential elements supposedly needed only in small quantities for healthy growth of plants (about 50-100 µM in the nutrient media). Under high external Cl^-^ condition, plant cells accumulate excessive Cl^-^ which causes toxicity. Thus, the chloride homeostasis in plant should be finely regulated. To further test if BIRG1-mediated chloride efflux affects chloride homeostasis in plant, we measured the chloride contents in WT and *birg1* mutant. As shown in Fig. **7**, the Cl^-^ content in the *birg1* mutant grains was obviously higher than that in the WT grains when grown under normal paddy field (Fig. **7a**), suggesting that BIRG1 regulates chloride homeostasis in the grain and the chloride content in the grain is positively correlated to the grain size. Furthermore, the Cl^-^ contents in the plants grown on 1/2MS medium and 1/2MS supplemented with 100 mM NaCl for 14 days were also determined. When grown on 1/2MS, the Cl^-^ content in the roots of WT and *birg1* mutant was comparable. The same is true in shoots (Fig. **7b, c**). When grown on 1/2MS supplemented with 100 mM NaCl, WT root and shoot accumulated more Cl^-^ compared with those grown on 1/2MS (Fig. **7b, c, d, e**). In *birg1* mutant, the Cl^-^ content in shoot was comparable to that in WT (Fig. **7e**). However, the *birg1* roots contain significantly more Cl^-^ than WT roots (Fig. **7d**). These results indicated that, in addition to its role in controlling chloride homeostasis in grain under normal growth condition, BIRG1 is essential for the chloride homeostasis in root under high salinity condition.

**Fig. 7.**
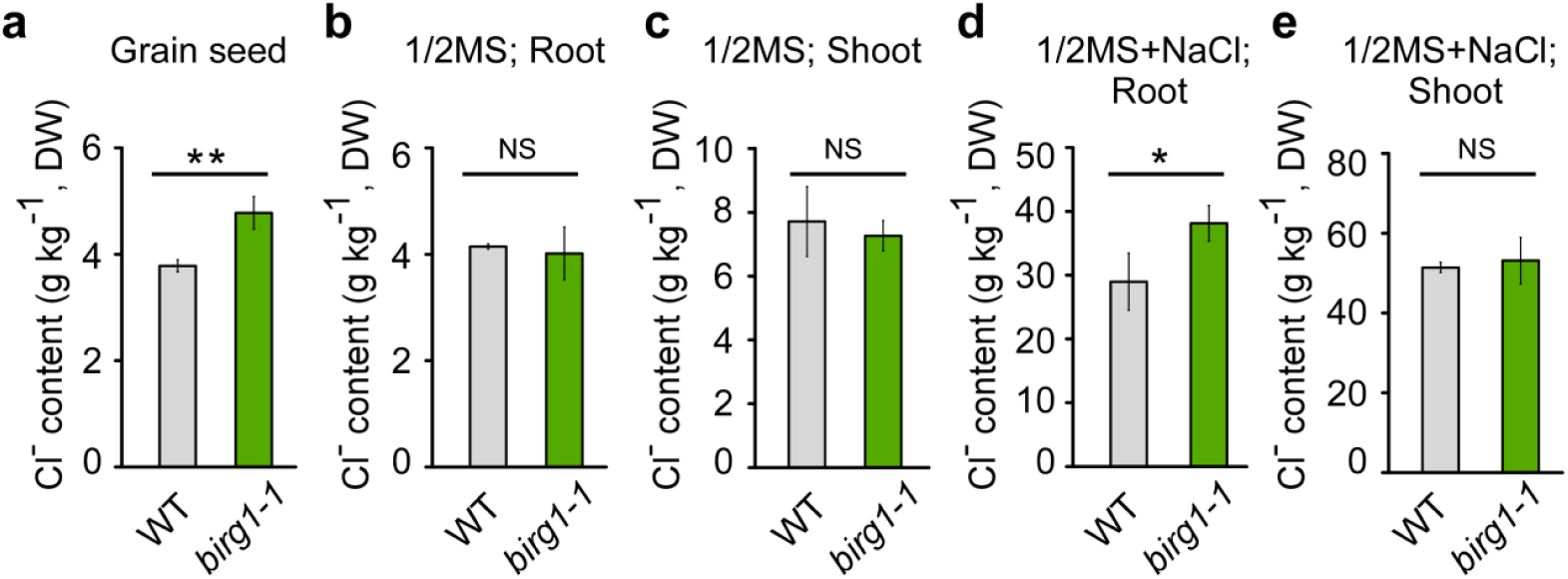
Chloride content of WT and *birg1* mutant. (a) Chloride content of WT and *birg1* grain. The rice plants were grown in a paddy field located in Lang Fang, He Bei, China, n=3. (b and c) Chloride content of the seedling root and shoot of WT and *birg1* grown on 1/2MS. (d and e) Chloride content of the seedling root and shoot of WT and *birg1* grown on 1/2MS plus 100 mM NaCl. 14-day-old seedling root and shoot were separated, and 6 roots or shoots were collected as one sample, n=3. The data are mean ± SD, Student’s *t*-test was used to generate the *p* values, * *p*<0.05, ** *p* <0.01 and *** *p* <0.001.

## Discussion

In this study, we demonstrate that a member of DTX/MATE family transporters, BIRG1, negatively regulates grain size by specific controlling cell size but not cell number of the spikelet hull. BIRG1 localizes to the PM and mainly expressed in reproductive organs and roots. BIRG1 shows Cl^-^ efflux activity in oocyte. In line with the transport activity of BIRG1, the *birg1* mutant is less tolerant to high external Cl^-^. The *birg1* grains contain higher level of chloride compared to WT grains when grown under normal paddy field and the *birg1* roots accumulate more chloride than those of WT under high salinity. Thus, we conclude that BIRG1 functions as a Cl^-^ effluxer that regulates grain size and salt tolerance by controlling chloride homeostasis in rice.

Grain size is one of the key agronomic traits that determine grain yield, but the genetic and molecular mechanisms of grain size control in rice are still unclear. Our finding that BIRG1 negatively regulates grain size and the *birg1* mutant grain contains more Cl^-^ than that of WT indicates that Cl^-^ content is positively correlated to the grain size under normal growth condition. It was demonstrated that the larger spikelet in *birg1* mutant resulted from the enlargement of the outer parenchyma cells (Fig. **4f, i**). Meanwhile, some of the cell expansion-related genes were significantly elevated in young panicles of the *birg1* mutant (Fig. **S7c, d**). Previous studies reported that Cl^-^ stimulates the activity of the tonoplast-type H^+^-translocating ATPase (Churchill and Sze, 1984; Randall and Sze, 1986), which contributes to the intrusion of anions (e.g. nitrate and chloride) into the vacuole. The anions, together with their counterparts, serve as osmotica in the vacuole, driving the water flow and enhancing cell turgor pressure. In this way, Cl^-^ is thought to promote turgor-driven cell expansion (Chen *et al*., 2016; Franco-Navarro *et al*., 2016; L.B. Smart *et al*., 1998). This may explain why the outer parenchyma cells of *birg1* mutant grain are enlarged. At the meantime, turgor pressure may serve as a signal that activates the expression of cell expansion-related genes to facilitate cell expansion (Beauzamy *et al*., 2014; Fricke *et al*., 2000).

We also noticed that under normal growth condition, the yield of *birg1* mutant is slightly lower than that of WT (Fig **S4m**), indicating a complicated physiological effect of Cl^-^ homeostasis in regulating agronomic traits of rice. Nevertheless, it will be worthwhile to figure out how BIRG1 is connected to the molecular pathway of grain size and yield regulation in the future.

NaCl is generally the dominant salt in saline soils. Both Na^+^ and Cl^-^ cause detrimental effect on plant growth and development if accumulated at high concentrations in the cytoplasm. However, most previous researches on salt tolerance have been focusing on Na^+^. Thus, it is still elusive that how Cl^-^ is transported and how Cl^-^ homeostasis is regulated under salt stress (Teakle and Tyerman, 2010). Our finding that BIRG1 mediates Cl^-^ exclusion in the root thereby contributing to the salt tolerance advances our understanding of Cl^-^ transport mechanisms under salt stress. More importantly, it is suggested that BIRG1 may server as a molecular target for engineering rice with improved salt tolerance.

Organic compounds, including a number of therapeutic drugs, have been shown to be substrates of DTX/MATE transporters in bacteria and animals (Shiomi *et al*., 1995; Morita, 1998). The DTX /MATE proteins in plants constitute a superfamily, and some of them have been shown to transport organic acids or secondary metabolites (Li *et al*., 2002; Magalhaes *et al*., 2007; Maron *et al*., 2009; Yokosho *et al*., 2009; Zhang *et al*., 2014; Kengo *et al*., 2016). For example, MATE proteins play a role in aluminum tolerance by mediating citric acid efflux from root cells to chelate Al^3+^ (Yokosho et al., 2011). Recently, two tonoplast MATE/DTX proteins, DTX33 and DTX35, have been demonstrated to function as turgor-regulating chloride channels in *Arabidopsis* (Zhang *et al*., 2017), expanding the diversity of substrates for plant DTX /MATE. Our finding that rice BIRG1 possess a strong Cl^-^ efflux activity that provides the first example in monocots and further solidifies the notion that some plant MATE/DTX proteins are halogen ion/anion channels, which breaks the dogma that DTX/MATE proteins are transporters of organic compounds.

## Methods

### Plant Materials and Growth Conditions

The *japonica* Nipponbare, as WT for analyses, was kindly provided by prof. Lu Tiegang. The rice plants were grown in paddy fields located in different regions of China, under natural conditions or on 1/2 MS (Murashige and Skoog) + 1% Sucose + 0.6% Agar (pH5.8) in a growth chamber (26-28°C, 16-h light/8-h dark). The mutants containing *birg1-1*, *birg1-2*, RNAi lines and other *dtxs* were achieved by using either TALENs or CRISPR/Cas9 technologies in the Nipponbare background. TALENs-*birg1-1* was designed and operated by prof. Gao Caixia’Lab. CRISPR/Cas9 lines were achieved from Beijing Genovo company.

### RNA extraction and Real-Time qRT-PCR Analysis

Total RNA was isolated from the various organs of *birg1* and wild-type plants using Trizol reagent (Life Technologies). About 1 μg RNA was reversely transcribed using oligo (dT) primer and was reverse-transcribed with Superscript III following the manufacturer’s instructions (Life Technologies). Quantitative PCR experiments were performed using SYBR Green PCR Master Mix (Takara, Otsu, Japan) with Bio-Rad CFX96 real-time PCR machine. The primers are listed in Supporting Information table 2. The rice *ACTIN1* gene was used as internal control. Values are means ±SD of three biological repeats. Changes in transcription were calculated using the 2 (-Delta Delta C (T)) method (Livak and Schmittgen, 2001). The primers are described in Supporting Information table 2.

### Vector Construction and Plant Transformation

For the subcellular localization assay, complementary cDNAs of *BIRG1* was inserted into *p*E3242-GFP to produce the 35S:: *BIRG1*-*GFP* constructs. Upon sequence confirmation, *BIRG1* full-length coding sequences were cloned into *p*CAMBIA1302 to generate the *p*CAMBIA1302-35S: *BIRG1* for tobacco epidermal leaf cells transiently transformation (Waadt and Kudla, 2008; Waadt *et al*., 2008). For the GUS-staining assay, a 3.2-kb native promoter was cloned into *p*CAMBIA1300-GN vector to generate *p*CAMBIA1300GN-*PRO BIRG1*::*GUS* construct. To generate *BIRG1*-RNAi construct, a 300-bp gene-specific fragment of *BIRG1* coding sequence was amplified and cloned into *p*TCK309 vector. All recombinant constructs were transformed into the rice genome by agrobacterium tumefaciens mediated transformation methods as described previously (Komari, 2010). The primer sequences for vector construction are listed in Supporting Information table 2.

### GUS staining and scanning electronic microscopy observation

GUS staining activity was detected according to the method as previously described (Jefferson and R., 1989). The samples were treated in GUS staining buffer overnight at 37°C. The samples were then cleared in 70% - 80% - 90% - 100% ethanol to remove chlorophy ll. The observed staining patterns were recorded with microscope (leica and Zeiss Stemi SV11). The spikelet hulls and dried seeds were prepared for scanning electron microscopy (HITACHI, S-3000N) observation following previous description (Jin *et al*., 2011).

### Measurement of grain filling rate

Representative samples of 200 grains from the main panicle at 2, 3, 6, 9, 12, 15, 18, 21, 24, 27 and 30 d after fertilization (DAF) were harvested. Half of them were used to measure grain fresh and dry weight. And the other half were carefully dehusked by hand and used to measure brown rice fresh and dry weight. Every 20 grains or brown rice as one point sample and then calculate its average.

### Histological analysis

Young spikelet hulls were fixed in FAA (5% glacial acetic acid: 5% formaldehyde: 50% ethanol=1:1:18) for 48 h, and then dehydrated in a graded ethanol series (70%, 80%, 90%, 100%, and 100%). Fixed tissues were embedded in Paraplast Plus chips (Sigma) and were cut into 10 μm thick sections using a microtome, then gradually rehydrated and dried before toluidine blue staining for light microscopy.

### Subcellular localization

The expression constructs 35S:: *BIRG1*-*GFP* and 35S::*GFP* were transformed into rice protoplasts by PEG-mediated transfection. And transfection of rice protoplasts were performed according to the method described previously with minor modifications (Chen *et al*., 2006; Yoo *et al*., 2007; Zhang *et al*., 2011). The *p*CAMBIA1302-35S:: *BIRG1* construct was transformated into tobacco epidermal leaf cells by agrobacterium-mediated transient expression in *N. benthamiana* as described previously (Olivier *et al*., 2003; Waadt and Kudla, 2008; Waadt *et al*., 2008). Confocal images were captured using Zeiss Live 780 equipment. GFP fluorescence was excited at 488 nm wavelength, and the emission filters were 500-530 nm.

### Two-Electrode Voltage-Clamp (TEVC) recording from *Xenopus* oocytes

The *BIRG1* cDNA was cloned into the *p*GEMHE oocyte expression vector. The capped cRNA was synthesised with the Ambion mMESSAGE mMACHINE T7 Kit. Each oocytes were injected with 32 nl of the cRNA in 500 ng μl^-1^ or with 32 nL of RNase-free water (control oocytes) for voltage-clamp recordings. After 3 days incubation in ND96 solution (96 mM NaCl, 2 mM KCl, 1 mM MgCl_2_, 1.8 mM CaCl_2_, 10 mM HEPES/NaOH, pH7.4) supplemented with 25 g L^-1^ gentamycin at 16-18°C, oocytes were voltage-clamped using a TEV 200A amplifier (Dagan Corporation) and monitored by computer through Digidata 1440A/D converter and pCLAMP10.4 software (Axon Instruments) as described previously (Liu and Luan, 2001; Tian *et al*., 2015) with some modifications. All electrodes were filled with 3 M KCl. The bath solutions contained a background of 2 mM K-gluconate, 1 mM Mg-gluconate_2_, 1.8 mM Ca-gluconate_2_ and 10 mM MES/Tris (pH5.6). The bath solutions added with 48 mM Cl^-^, NO_3_^-^, SO_4_^2-^, Malate^2-^, Glu^-^ and Br^-^ were used for ion-selective analysis. Different concentrations of Na-Glu (24, 48, 96 mM) were added to the bath solution for concentration dependence experiments. Perfusion bath solutions contain 48 mM Cl^-^ along with 0, 10 or 100 mM DIDS were used for pharmacological analysis. D-Mannitol was added to adjust the osmolarity to 220 mOsmol kg^-1^. Voltage steps were applied from +40 to -140 mV in -10 mV decrements during 2.0 s. And the holding potential was seted to 0 mV.

### Determination of Rice Grain Quality

The protein content was measured by the dumas principle method (Kjeltec System 1002, Tecator). The contents of amylose were measured by Double Wavelength colorimetry (Song *et al*., 2007). Determination gelatinization poperites in cereal and starch by viscograph. All assays were measured in State Administration of Grain.

### Cl^-^ Content Measurement

Every grain sample contains 20 g muture grain of WT and *birg1* mutant. Seedlings were grown on 1/2MS or 1/2MS +100 mM NaCl for 14 days. Six seedling roots or shoots formed one sample. Before the whole plants were harvested and used for measuring Cl^-^ contents, they should be rinsed at least three times with distilled water. Grain, seedling root or shoot samples were dried at 80°C for 2 days. And then grinded to powder. Cl^-^ contents were determined by Ion Chromatography (Thermo ICS-600). Three independent experiments were carried out. The soil Cl^-^ contents were determined by a modified silver titration method that has been described previously (Li *et al*., 2008b).

### The photosynthetic capacity measurement

The photosynthetic capacity recordings from mature flag leaves of 2-month-old plants were conducted using a Li-6400 infrared (IRGA)-based gas-exchange analyzer with a fluorometer chamber (Li-Cor Inc.) (Li *et al*., 2016).

## Author Contributions

L.L., C.H., W.T., Z.R. conceived and designed the project. Z.R. and C.H. performed molecular cloning and transgenic plant generation; Z.R. conducted the electrophysiological experiments; C.H., F.B. and C.F. conducted subcellular localization in protoplasts; J.X., L.W., X.W. and Z.R. conducted Promoter-GUS experiments and field experiments; M.L., J.X. and J.Q. conducted TALEN experiments; Q.Z., J.S., Q.N performed RNA extraction, cDNA synthesis, and qRT-PCR analysis; All experiments were independently reproduced in the laboratory. Z.R., W.T. and L.L. analyzed the data and wrote the manuscript. All authors discussed the results and commented on the manuscript.

## Acknowledgments

This work was supported by grants from the National Key Research and Development Program of China (YFD0300102-3 to L.G.L.), the General Program of National Natural Science Foundation of China (No. 31872170 to L.L. and No. 31900234 to C. H.), the Key Program of the National Natural Science Foundation of China (31930010 to L. L) and the Capacity Building for Sci-Tech Innovation-Fundamental Scientific Research Funds (19530050165 to L. L.).

## Supporting Information

**Fig. S1.**
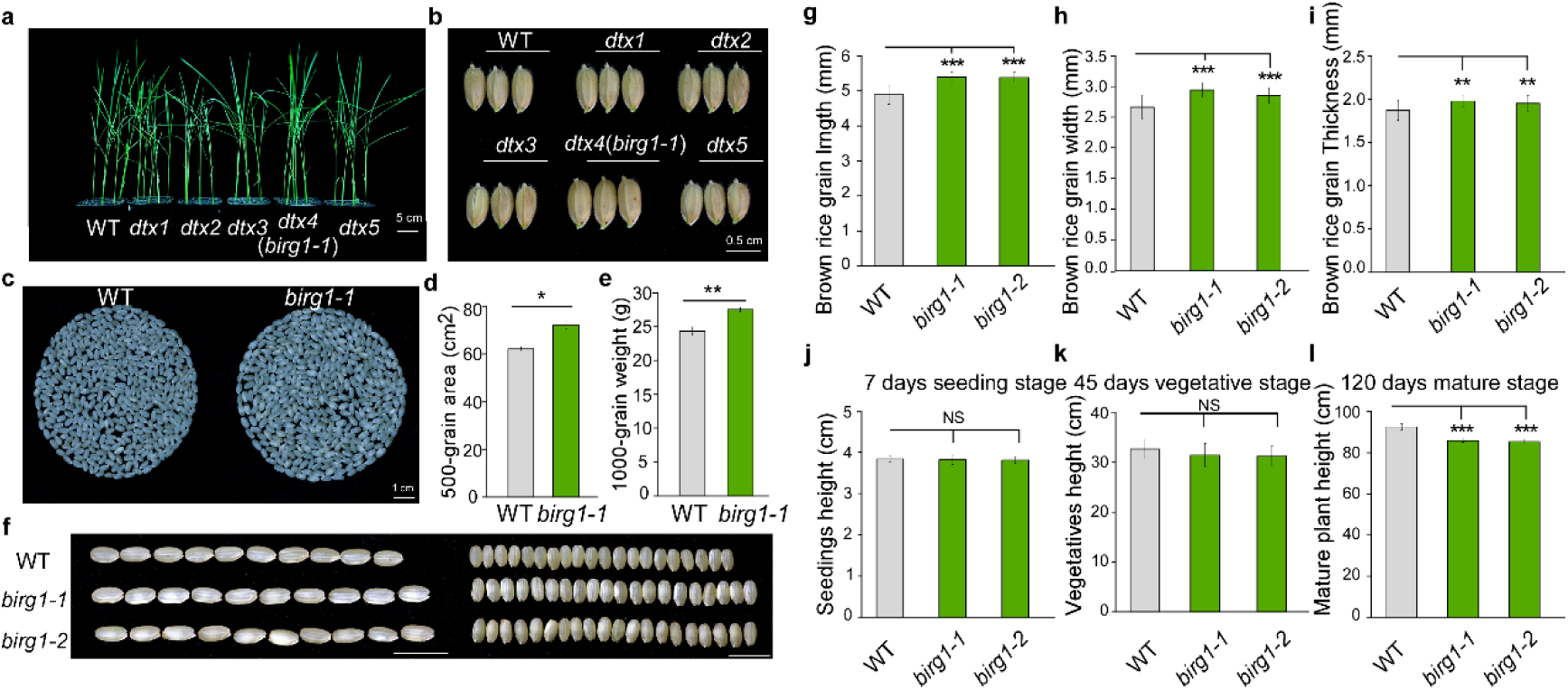
Phenotypes of *dtx* mutants. (a) Seeding morphology of the WT and several *dtx* mutants. The *dtx4* is named *birg1-1*. (Scale bar=5 cm). (b) Grain size of the WT and five *dtx* mutants. (Scale bar=0.5 cm). (c) Cover area of 500-grain in the WT and *birg1-1*. (Scale bar=1cm). (d) Statistical area of 500-grain in the WT and *birg1-1*. (e) Statistical data of 1,000-grain weight in the WT and *birg1-1*. (f) Brown rice morphology of the WT, *birg1-1* and *birg1-2*. Scale bars=1 cm. (g) Statistical data of grain length (j), grain width (h) and grain thickness (i) of the WT, *birg1-1* and *birg1-2.* n= 20. (j) 7 days seeding height of the WT and *birg1-1*, *birg1-2*. n=5. (k) 45 days vegetatives stage height of WT, *birg1-1* and *birg1-2*, n=8. (l) 120 days mature stage height of the WT and *birg1-1*, *birg1-2*. n=5. Data shown as means ± SD, *p* Value based on Student’s *t*-test, * *p*<0.05, ** *p* <0.01 and *** *p* <0.001.

**Fig. S2.**
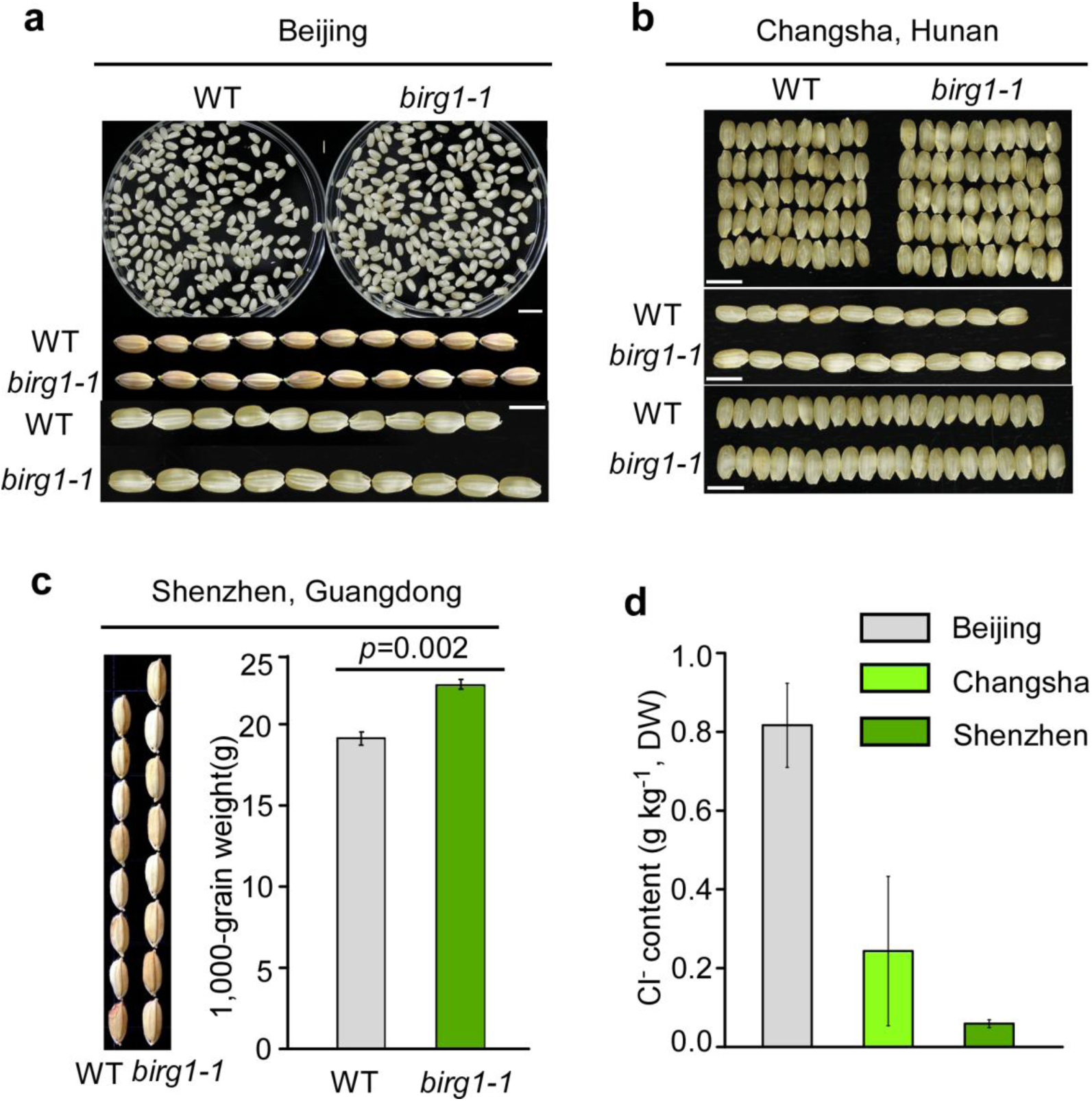
Grain phenotype of WT and *birg1* mutant in different regions with varied Cl^-^ level. Grain size of WT and *birg1* grown in Beijing, the north of China (a), (Scale bar=1 cm, 0.5 cm); Changsha, the middle of China (b) (Scale bar=0.5 cm); or Shenzhen, the south of China (c). The Cl^-^ contents in soil from Beijing, Changsha and Shenzhen were measured (d), n>=3. Data are represented as mean ± SD. *p* Value based on Student’s *t*-test.

**Fig. S3.**
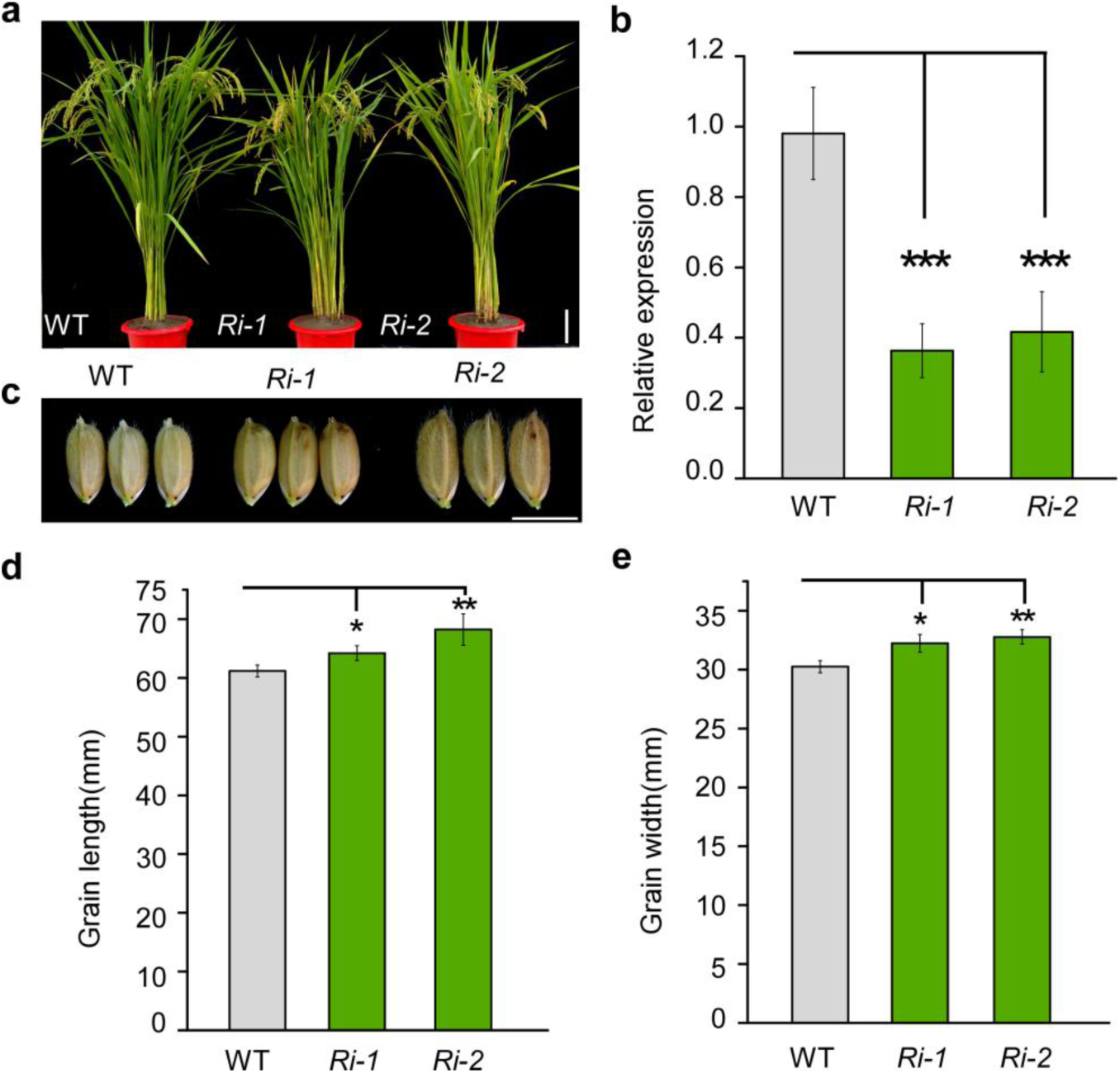
The phenotypes of *BIRG1*-knockdown plants (*Ris*). (a) The morphology of the WT and *Ris* (Scale bars=5 cm). (b) The expression levels of *BIRG1* in the WT, *Ri-1* and *Ri-2* plants detected by qRT-PCR. (c) Grain size of the WT, *Ri-1*and *Ri-2* plants (Scale bars=0.5 cm). (d) Statistical data of the grain length of the WT, *Ri-1* and *Ri-2*. (e) Statistical data of the grain width of the WT, *Ri-1* and *Ri-2*, Means ± SD are given in D and E (n =10), *P* value (Student’s *t*-test), * *p*<0.05, ** *p* <0.01 and *** *p* <0.001.

**Fig. S4.**
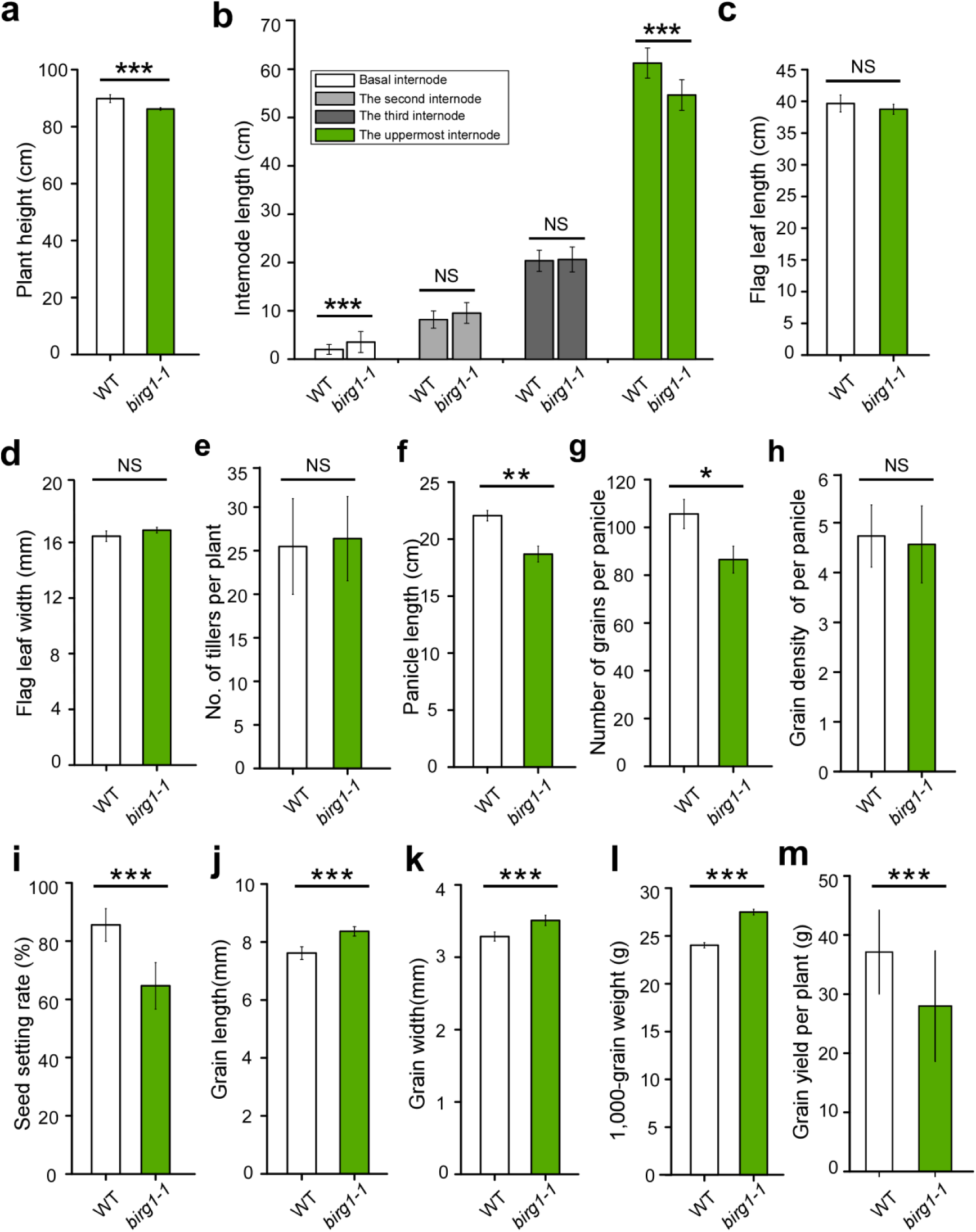
Agronomic traits of the WT and *birg1*. (a) Plant height of the WT and *birg1* as in (A), n=20. (b) Flag leaf length from different plants, n=20. (c) Flag leaf width (Measure the widest area), n=20. (d) Internode length, n=20. (e) Tiller number (Effective tiller number), n=20. (f) Panicle length, n=20. (g) Number of grains per panicle, n=20. (h) Grain density of per panicle, n=20. (i) Seed setting rate, n=20. (j) Grain length, n=20. (k) Grain width, n=20. (l) 1,000-grain weight, n=3. (m) Grain yield per plant, n=30. Data shown as means ± SD, *p* Value based on Student’s *t*-test, * *p*<0.05, ** *p* <0.01 and *** *p* <0.001.

**Fig. S5.**
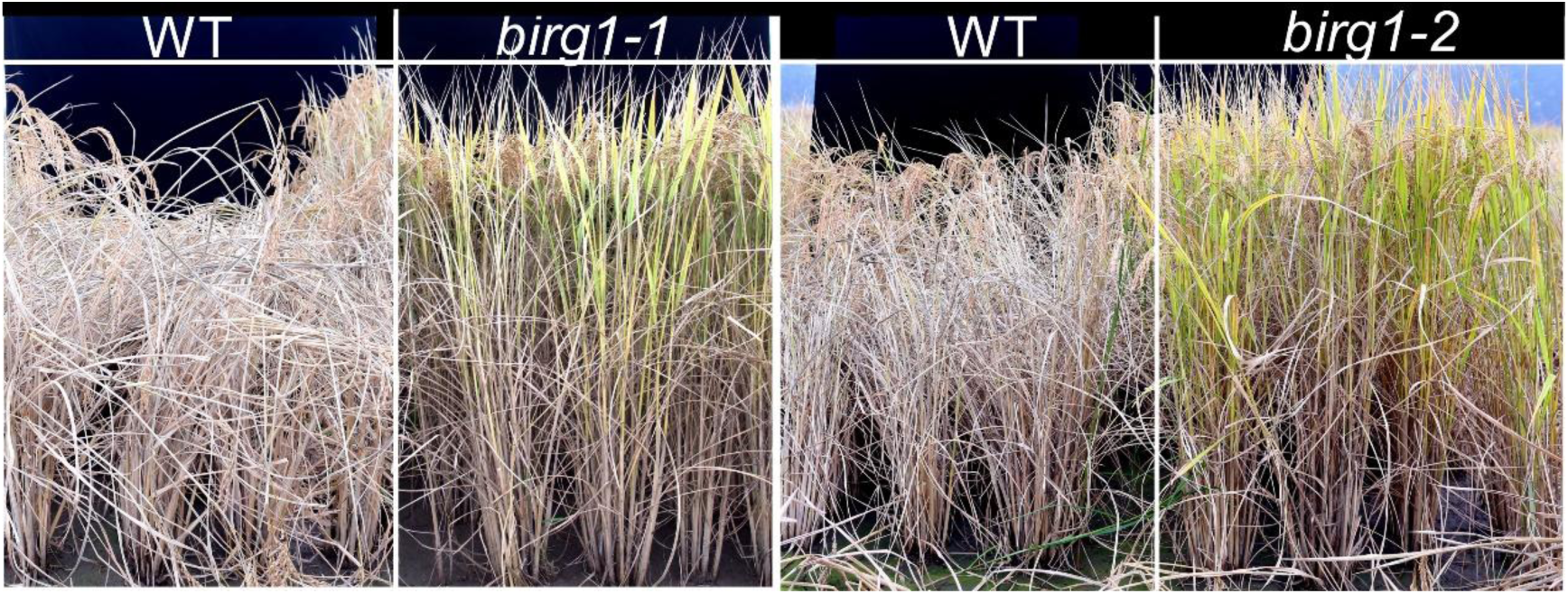
The *birg1* mutants exhibit improved lodging resistance. WT and *birg1* at the mature stage were shown.

**Fig. S6.**
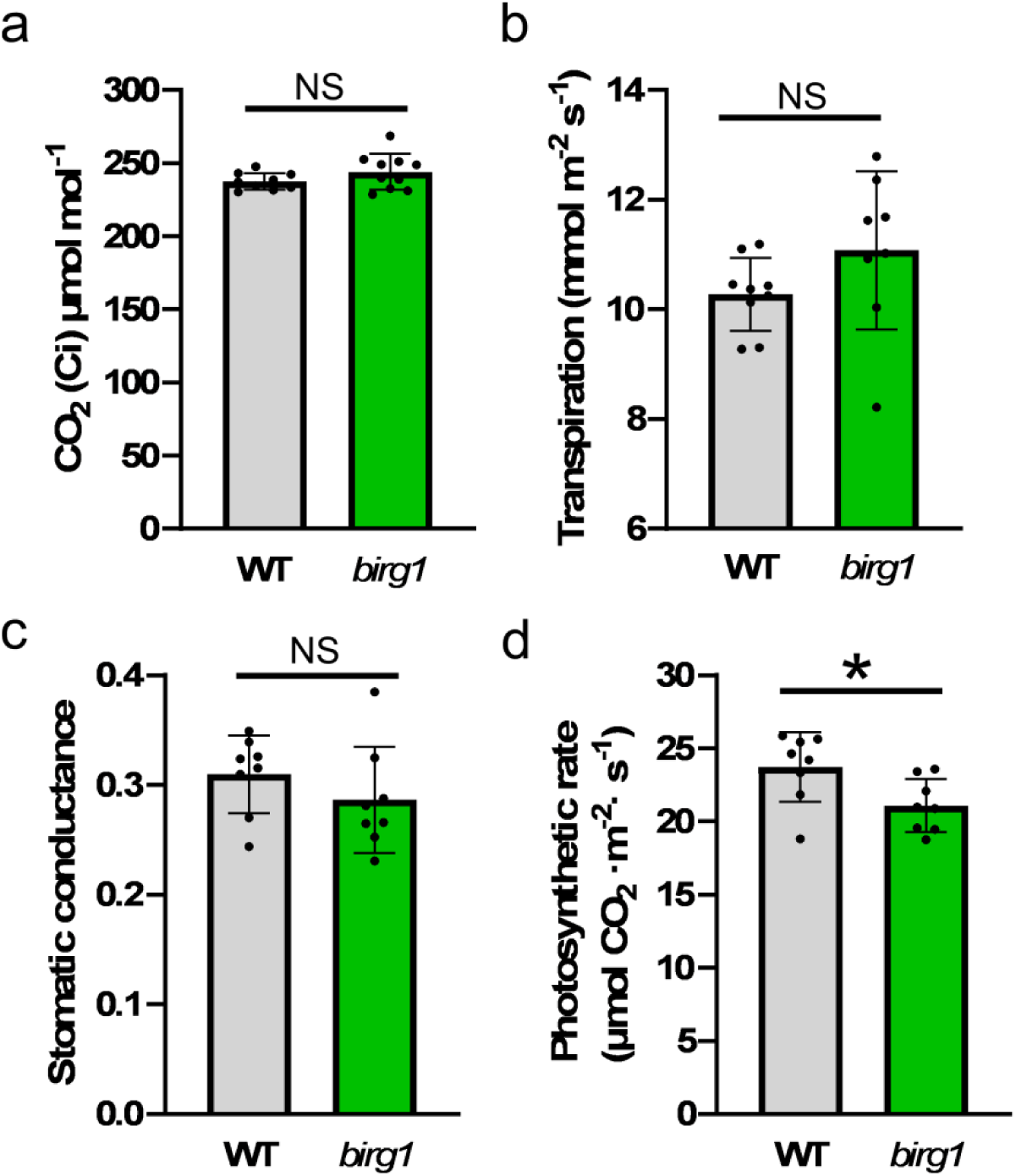
The photosynthetic capacity of WT and *birg1*. (a) Intercellular CO_2_ concentration. (b) Transpiration efficiency. (c) Stomatic conductance. (d) photosynthetic rate. Means ± SD are given (n >=8), *p* value (Student’s *t*-test, two-tailed), * *p*<0.05.

**Fig. S7.**
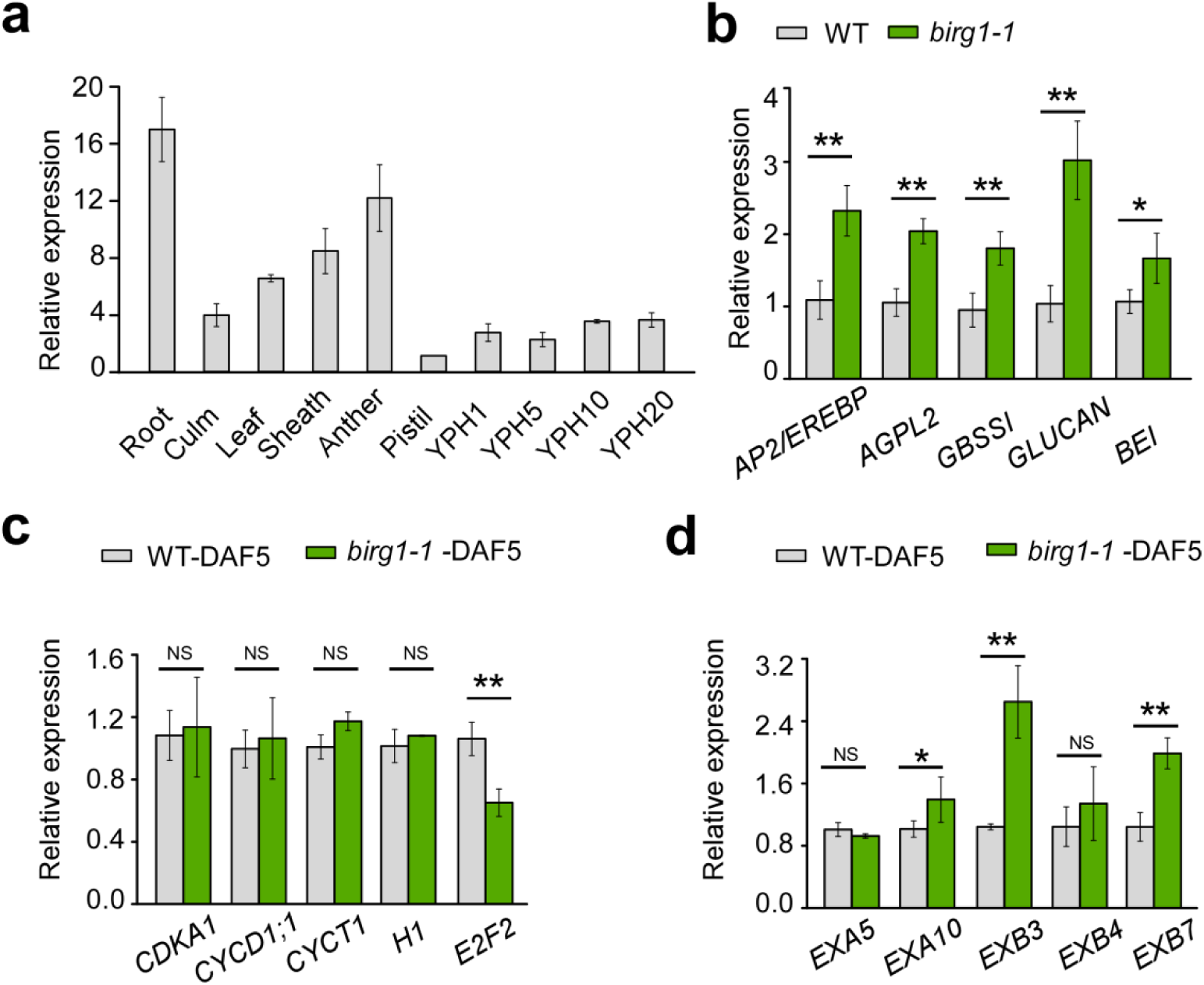
Expression of *BIRG1* in various organs and expression of genes involved in starch synthesis, cell cycle and cell expansion related genes in the WT and *birg1* analyzed by QRT-PCR. (a) Root, culm, leaf, and sheath were harvested from before heading WT plants. Anther and pistil were collected from spikelet hulls two days before fertilization. Young panicles heading (YPH) in different lengths (indicated as numbers, cm) were included for the analysis. (b) QRT-PCR analysis showed that expression levels of starch synthesis-related genes in *birg1* mutant were elevated compared with that in WT. Starch synthesis-related genes and ID: *RSR1/AP2/EREBP* (*Os05g0121600*), *AGPL2* (*Os01g0633100*), *GBSSI* (*Os06g0133000*), *GLUCAN* (*Os01g0533100*), *BEI* (*Os06g0726400*). (c)Transcription levels of cell cycle (G1 to S phase)-related genes in 5 DAF seeds of *birg1* relative to that of the WT. Cell cycle-related genes: *CDKA1* (*Os03g0118400*), *CYCD1;1* (*Os06g0236600*), *CYCT1* (*Os02g0133000*), *H1* (*Os03g0737600*) and *E2F2* (*Os12g0158800*). (d) Transcription levels of cell expansion-related genes in 5 DAF seeds of *birg1* relative to that of the WT. Cell expansion-related genes: *EXA5* (*Os02g0744200*), *EXA10* (*Os04g0583500*), *EXB3* (*Os10g0555900*), *EXPB4* (*Os10g0556100*) and *EXB7 (Os03g0102700)*. The rice *ACTIN1* gene was used as an internal control. Error bars shown as mens ± SD, and Student’s *t*-test identify significant differences at the *p* < 0.05, * *p*<0.05, ** *p* <0.01 and *** *p* <0.001.

**Fig. S8.**
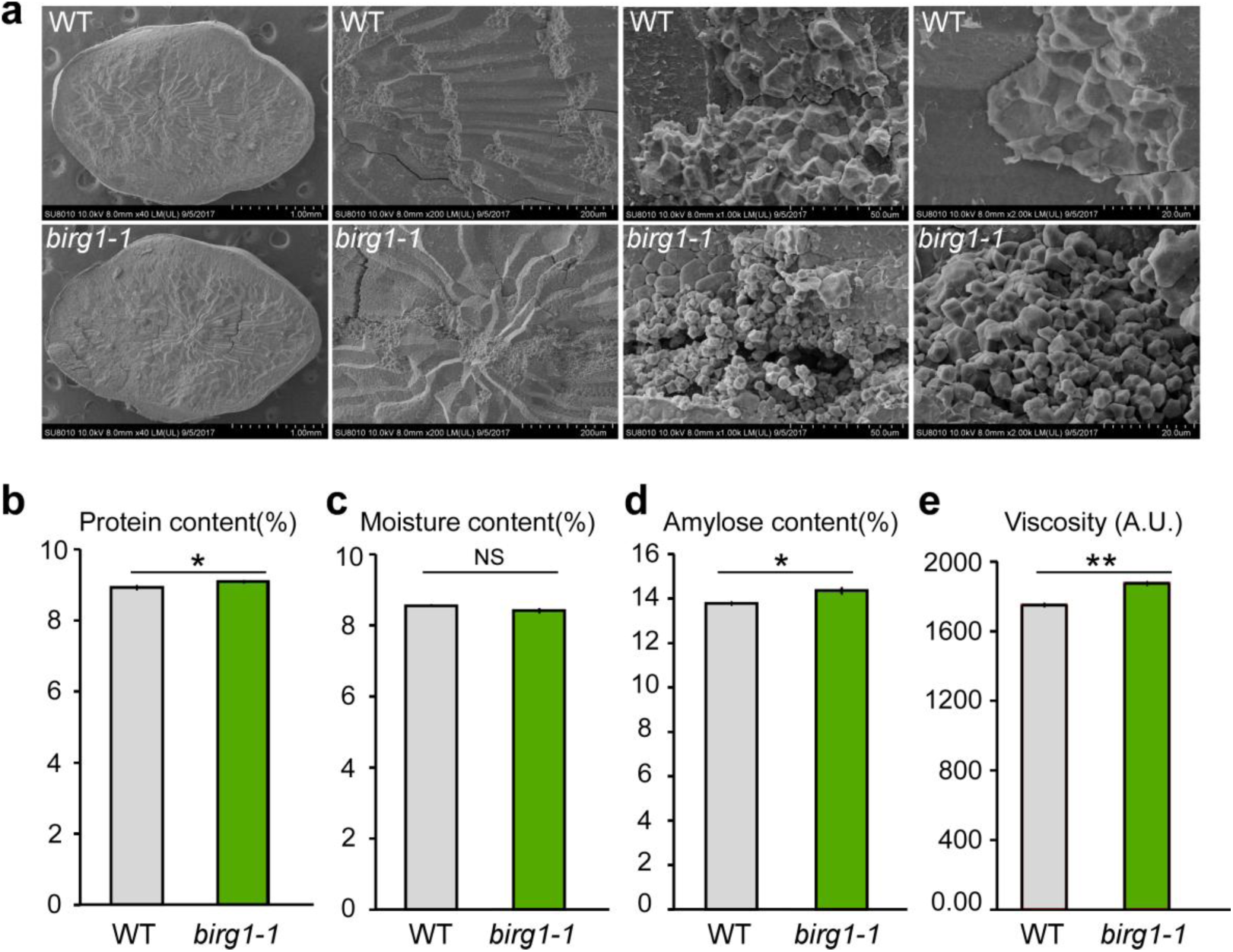
Scanning electron microscopy (SEM) analysis of WT and *birg1* mutant endosperm and physicochemical properties of starch in *birg1* grain. (a) The *birg1* mutant showed smaller and loosely packed starch grains as compared to large and tightly packed ones in the WT. Scale bars: 1 mm (the left row), 200 μm (second row), 50 μm (third row), 20 μm (right row). (b-d) Protein (b), moisture (c) and amylose (d) contents in mature caryopses of the WT and *birg1* mutant. (e) The viscosity of *birg1* pasting starch was higher than that of the WT. Data shown as mens ± SD (n = 3; \**p* <0.05, ** *p* <0.01, Student’s *t*-test).

**Table S1.** DTX/MATE family genes highly expressed in spikes

| Candidate genes | Gene NO. | TRAP-Seq FPKM Expression |
| --- | --- | --- |
| *OsDTX1* | *LOC_Os06g29950* | Seeding, Seed (S5 Stage) |
| *OsDTX2* | *LOC_Os01g49120* | Seeding, leaf, Seed (S4 and S5 Stage) |
| *OsDTX3* | *LOC_Os02g57570* | Seed (S3 and S4 Stage) |
| *OsDTX4* | *LOC_Os03g64150* | Root, Seed (S3 and S4 Stage) |
| *OsDTX5* | *LOC_Os08g43250* | Panicles, Seed (S4 Stage) |
| *OsDTX6* | *LOC_Os03g12790* | Seed (S4 and S5 Stage) |
| *OsDTX7* | *LOC_Os03g62270* | Root, Seeding |
| *OsDTX8* | *LOC_Os08g44870* | Seeding, leaf |
| *OsDTX9* | *LOC_Os11g03484* | Seed (S4 and S5 Stage) |
| *OsDTX10* | *LOC_Os08g43654* | Root, Seeding, Panicles |

**Table S2.**
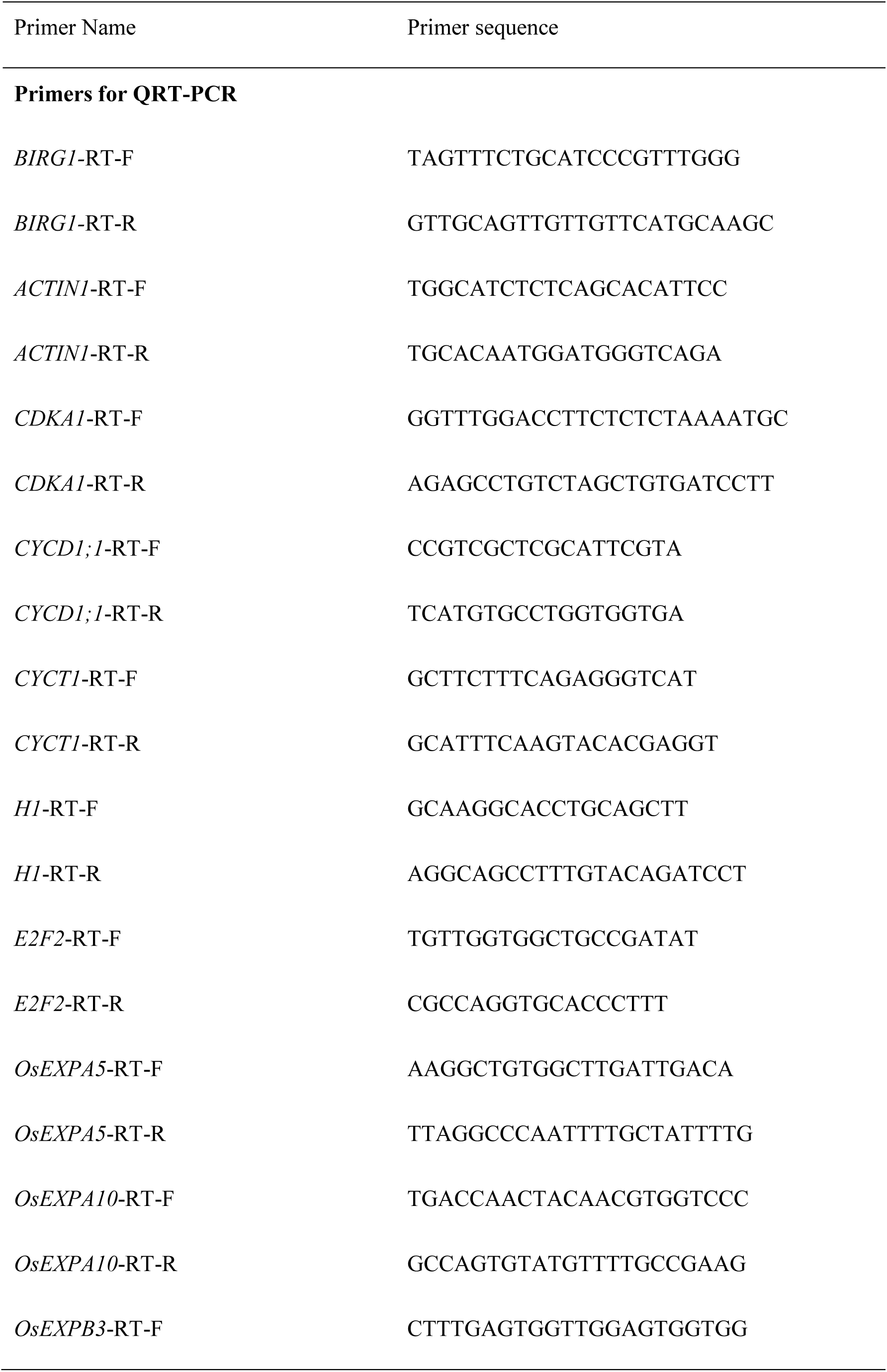
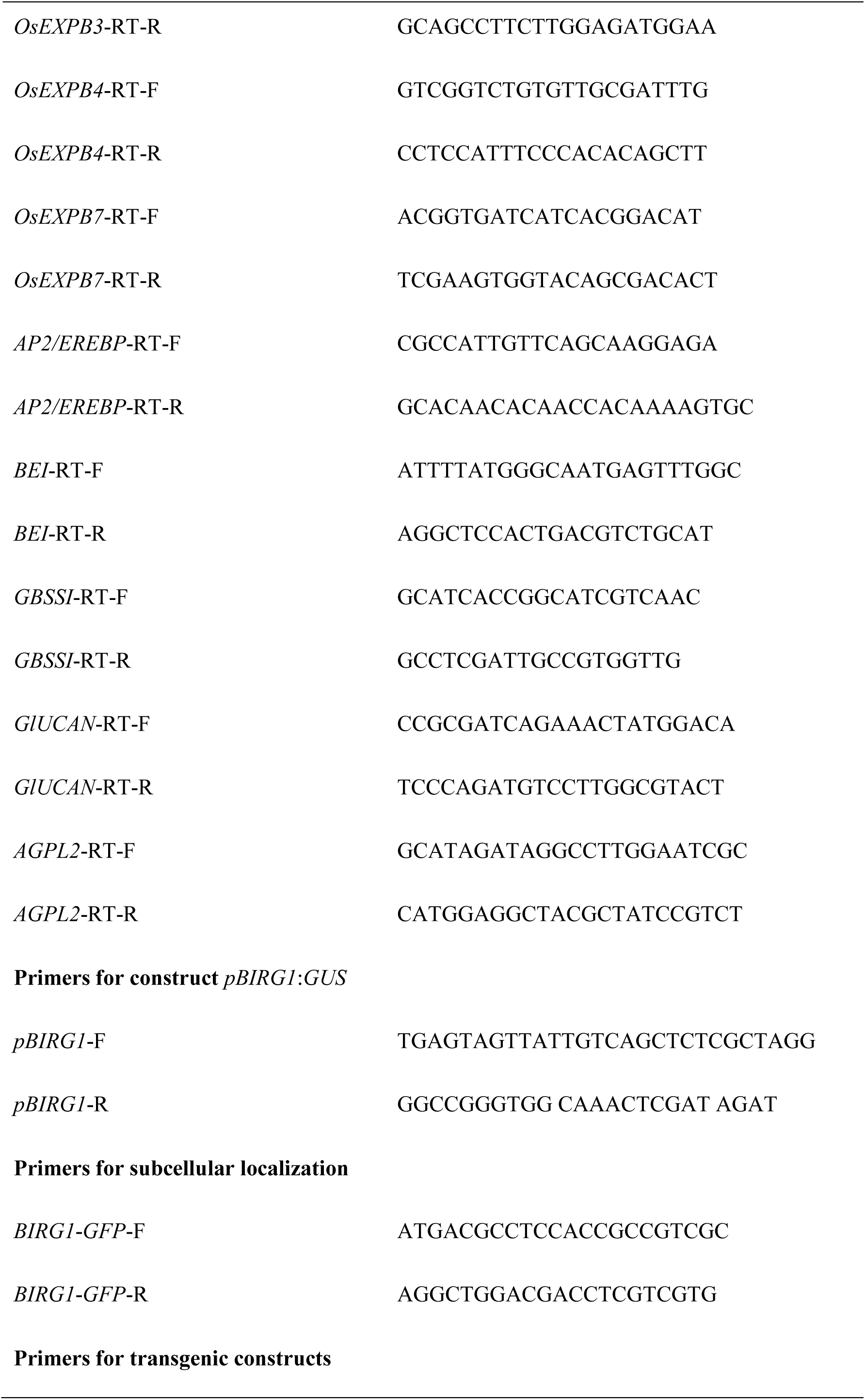
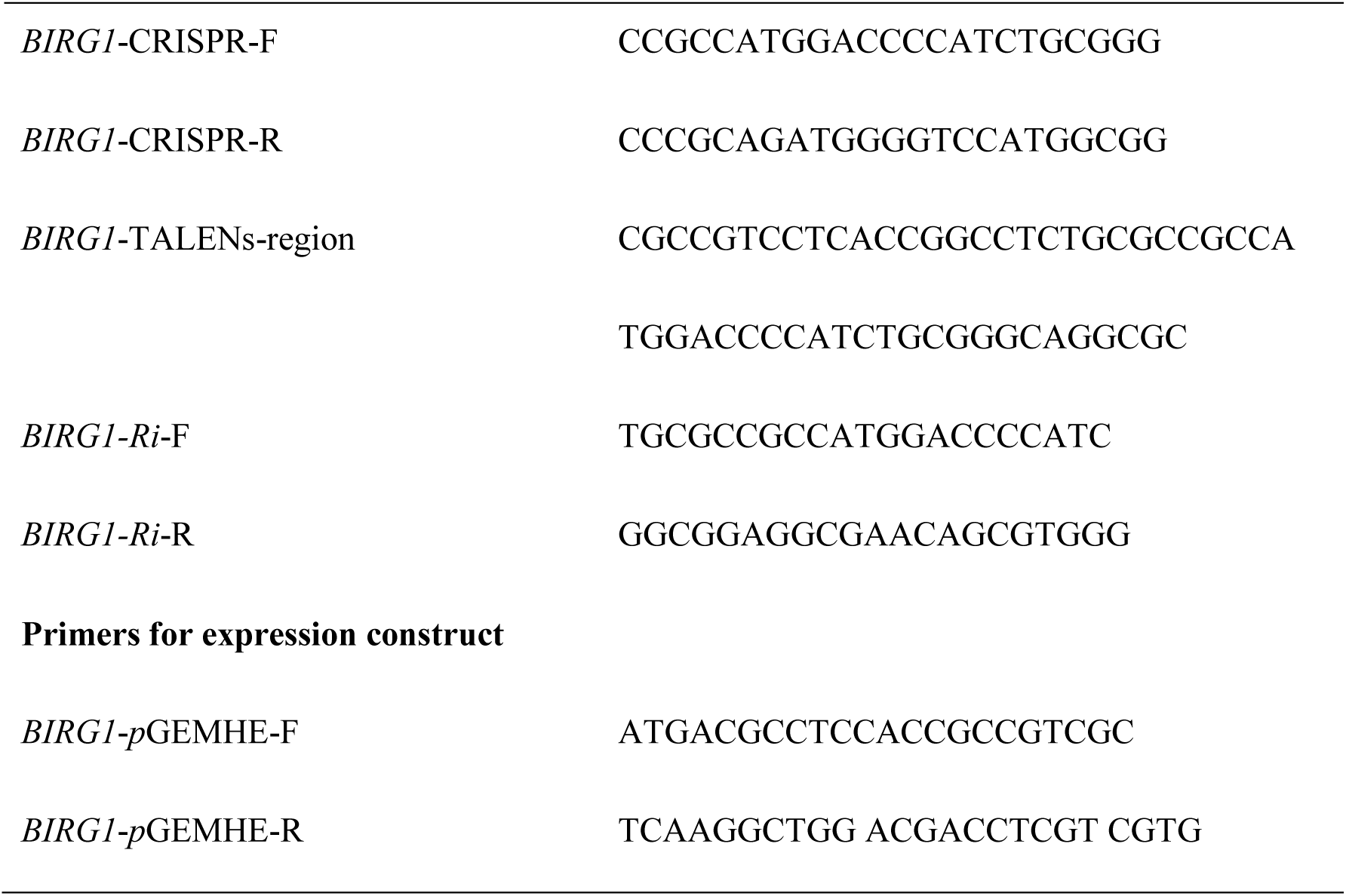
Primers used in this study

